# Volume regulation of cancer cells during osmotic pressure variation

**DOI:** 10.64898/2026.03.09.710325

**Authors:** Xiapeng Wang, Jiarui Gan, Wenjie Wu, Shubo Zhang, Ting Zhang, Chen Wang, Ye Chen, Qingchuan Zhang, Shangquan Wu

## Abstract

Osmotic pressure is a fundamental physical determinant of cellular homeostasis, and its perturbation is recognized as a critical factor in cancer progression and therapy. However, whether cancer cells and normal cells respond to osmotic pressure changes differently remains unclear. Here, we subjected cancer and normal cells to osmotic shocks and compared their volume and traction force changes. Under hypotonic conditions, cancer cells recovered their volume and forces much more slowly than normal cells, although both cell types responded similarly to hypertonic shock. We found that actin-based cortical tension controlled the recovery speed; whereas myosin-mediated contractility tuned the volume change extent. Substrate stiffness also influenced the recovery process by altering cytoskeletal constraint. A theoretical model integrating adhesion energy, ion and water transport, and surface tension was developed. The results matched the experimental observations. Our work uncovered the distinct mechanobiological behaviors of cancer cells upon hypotonic shock, providing a mechanical framework for understanding cancer mechanisms that may inform the development of cancer therapeutics.

## Introduction

Osmotic pressure is one of the most important physical factors that influence cell properties and activities, including cell growth[1] and differentiation[2]. When the extracellular osmotic pressure changes in a short period of time, the functions and activities of cells can be affected[3]. In tumor pathology, variations in osmotic pressure are closely connected to cancer progression[4] and therapy[5]. Studies have proved that asymmetric alterations in extracellular osmotic pressure can induce directional migration in cancer cells[6], providing new insights into the mechanisms of cancer cell invasion within confined spaces. In the context of cancer therapy, current clinical evidence indicates that establishing a hypotonic extracellular environment can provide adjuvant therapeutic benefits[7, 8]. In vitro studies have discovered that several cancer cell lines swell and then rupture after being treated with distilled water in 3-10 minutes, indicating the cytocidal effects of the extreme hypotonic stress on cancer cells[9].

Under steady-state conditions, water is in thermodynamic equilibrium across the plasma membrane, that is, the osmotic concentrations of the intracellular and extracellular compartments are nearly equilibrated. A change in solute concentration generates an osmotic gradient, thereby driving the immediate transmembrane flow of water to restore equilibrium[1, 10]. Given that animal cells cannot sustain substantial hydrostatic pressure differences, transmembrane water flows result directly in cell swelling or shrinkage[11]. To mitigate the detrimental effects caused by acute osmotic shocks within a short timeframe, cells respond to drastic volume changes by activating the volume regulation systems known as regulatory volume decrease or regulatory volume increase[12]. Cellular volume regulation relies on the transport of osmotically active solutes, mediated by the coordinated action of specific channels and exchangers including the Na^+^-K^+^-2Cl^−^ cotransporter (NKCC1), Na^+^/ H^+^ exchanger (NHE1) and Cl^−^/HCO3^−^ exchanger (AE2)[10, 13]. Studies on suspended cells under osmotic shock have provided a foundation for understanding the general cellular response patterns. The mechanisms of how the suspended cells respond, particularly the underlying volume regulation principles, have been studied by previous researchers[14, 15]. For instance, a theoretical model incorporating water permeation, mechanosensitive channels, active ion pumps, and active stresses in the cortex to describe how the volume and pressure of suspended cells are regulated would it differ through the process of osmotic shock has been well established[16]. Compared to suspended cells, adherent cells are connected to the extracellular matrix, so that their physical microenvironment more closely resembles the in vivo condition. Unlike suspended cells, substrate-attached cells experience a basal constraint, which potentially leads to different behavioral responses. For instance, the mechanical properties of the external substrate (e.g., stiffness) might impact cellular behaviors, one of which is the cellular behavior under osmotic shock[17-19]. Under osmotic shock, cells deform in response to the drastic change in extracellular osmolality. This deformation and the subsequent volume adjustment involve strain in the cortical layer. Therefore, differences in cortical mechanical properties and mechanical behaviors may lead to variations in the cells’ deformation response. Cells undergo alterations in their mechanical properties following the onset of disease. Notably, the mechanical characteristics of cancer cells differ significantly from those of normal cells[20, 21]. Thus, cancer cells and normal cells may exhibit distinct behavioral responses under osmotic shock. Hence, investigating whether the distinction exists and if so in what pattern it would differ may contribute to a deeper understanding of cancer pathogenesis and provide novel mechanical insights for designing targeted therapies to eliminate cancer cells.

In this study, we aim to reveal the volume and traction force responses of cancer cells and normal cells to osmotic shocks. Using pharmacological modulation of the cell cortex, we further investigate the resulting volume response and the underlying regulatory mechanisms. We also characterize the temporal dynamics of the cellular actin network in response to osmotic shock. Furthermore, we investigate how cells respond to osmotic shocks when the mechanical properties of the substrate change. Additionally, we propose a theoretical model that incorporates dynamic adhesion energy to describe shape regulation in adherent cells under osmotic shocks.

## Results

### Cancer and Normal Cells Volume Responding Distinctions

In order to investigate the volume response of cancer cells and normal cells under osmotic shocks, we measured volume changes in breast cancer cell line MDA-MB-231 and normal breast cell line MCF-10A exposed to osmotic shocks (Fig. 1A). Due to their weak adherence and hemispherical morphology, volume changes were reflected by variations in the projected cell area. To verify the validity of our approximation, we labeled the F-actin and captured real-time 3D images, which showed that the volume changes aligned with the projected area changes (Fig. S1 and S2). Upon hypotonic shock (from ∼ 300 mOsm to ∼ 100 mOsm), both types of cells exhibited immediate swelling, with their areas reaching the peak at around 100 seconds (Fig. 1A and B). The peak areas were about 1.7-fold of their original areas. During the swelling phase, cancer cells and normal cells demonstrated no significant difference in their expansion rate. After reaching the peak, there followed a regulatory volume decrease (RVD) phase, during which the two types of cells exhibited markedly different recovery kinetics: MDA-MB-231 cells recovered only about 20% within 200 seconds after shock, whereas MCF-10A cells recovered up to approximately 60% (Fig. 1B and C). At 500 seconds after hypotonic shock, normal cells had almost completely recovered their volume, whereas cancer cells showed minimal recovery. Under a milder hypotonic shock (200 mOsm), both cell lines reached a smaller peak of about 1.6-fold, and MDA-MB-231 cells still recovered volume significantly more slowly than MCF-10A cells (Fig. S3 A and B). This reproducible pattern across different osmotic gradients confirms the reliability of our findings that cancer cells have a notably slower volume recovery rate compared to normal cells under hypotonic shocks. To further investigate the response patterns of cells to osmotic shocks, we applied hypertonic shocks on both types of cells (Fig. 1A and B). Hypertonic shock (600 mOsm) induced by PBS of high osmolarity led to immediate shrinkage. Cell area decreased by about 25% in 60 seconds and barely any recovery followed in 500 seconds for both types of cells. We further used mannitol solution prepared in PBS (450 mOsm) to induce a milder hypertonic shock. In both cancer cells and normal cells, 20% area reduction and later a very slight regulatory volume increase were observed (Fig. S3C). Under hypertonic conditions, the volumetric responses of normal and cancer cells were strikingly similar. Both the extent and the rate of volume recovery of cancer cells and normal cells were nearly identical, in sharp contrast to the differential behavior observed under hypotonic shock. To investigate the universality of the differential volume recovery between cancer and normal cells under hypotonic shocks, we further tested liver cancer cell line HepG2 and normal liver cell line THLE-2 under 200 mOsm hypotonic shock (Fig. S3G). The volume expansion phases and peaks of both types of cells showed almost no difference and recovery rate of cancer cell HepG2 was evidently slower (Fig. S3 D and E), exhibiting a similar trend as the breast cell lines. All these results above demonstrate that under hypotonic shock, cancer cells and normal cells undergo the same expansion phase but their recovery dynamics differ substantially (Fig. 1D). Specifically, cancer cells exhibit evidently slower volume recovery rate. When under hypertonic shock, however, similar RVI kinetics were observed in both cell types.

**Fig. 1.**
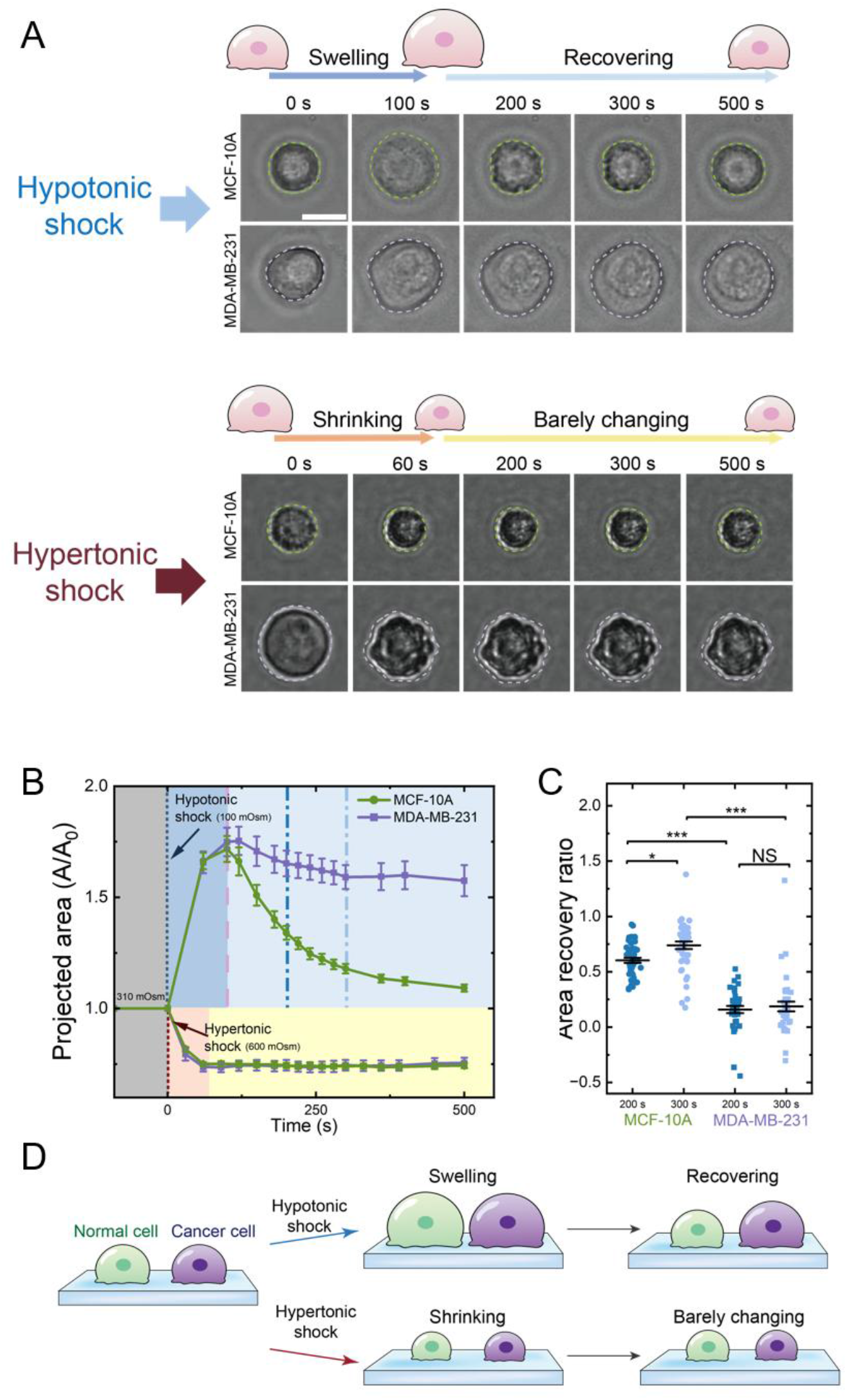
Volume changes under osmotic shocks. (**A**) Bright-field images of projected area changes of normal cell MCF-10A and cancer cell MDA-MB-231 (Scale bar: 10 μm). (**B**) Averaged normalized projected area change over time following osmotic shock (MCF-10A: 100 mOsm, n=14; 600 mOsm, n=10. MDA-MB-231: 100 mOsm, n=15; 600 mOsm, n=9; mean ± SE). (**C**) Area recovery ratio 200 s and 300 s after 100 mOsm hypotonic shock, calculated as (the maximum area - the recovery area)/(the maximum area – the initial area); Data are mean ± SE (MCF-10A, n = 45; MDA-MB-231, n = 37; mean ± SE). (**D**) Schematic summarizing the differential volume recovery dynamics between cancer and normal cells under osmotic shock.The figure legend should begin with a title (an overall description of the figure, in boldface) followed by additional text. Each legend should be placed immediately after its corresponding figure.

### Coordinated Traction-Volume Responses

Cells are connected to the underlying ECM via transmembrane proteins. Cellular deformations are therefore transmitted to the substrate, exerting forces and causing substrate deformations. We quantified the force changes by measuring the associated substrate displacements (Fig. 2A)[22]. The total traction was defined as the algebraic sum of traction forces over the cell-covered area, and the mean traction was obtained by averaging the values across all points. Prior to hypotonic shock, the initial mean traction force of MDA-MB-231 was higher than that of MCF-10A (Fig. 2A and Fig. S5B), as MDA-MB-231 cells exhibited more basal stress fibers (Fig. S7A)[23]. Upon hypotonic shock, the mean traction force dropped at a similar rate in both cell types, reaching a minimum around 100 seconds; however, the subsequent recovery patterns diverged significantly (Fig. 2A-C). For MDA-MB-231 cells, the traction force remained at this minimum in a sustained plateau, showing no recovery (Fig. 2A and C). In contrast, for MCF-10A cells, the minimum force marked the onset of recovery (Fig. 2A and B). The recovery lasted about 150 seconds, stabilizing at around 88% of the initial mean traction force. The traction force dynamics mirrored the volume changes (Fig. 2B and C), consistent with the established coupling between cortical tension and traction force[24]. Under hypotonic shock, the traction force dynamics in MCF-10A cells followed a pattern comparable to that reported for C2C12 cells[24]: an initial drop during volume expansion followed by recovery during volume regulation. In contrast, traction force in cancer cells declined without subsequent recovery. This indicated that differences in volume regulation directly led to corresponding variations in traction force response (Fig. 2D). As traction force reflects cell-ECM interactions, these differing dynamics indicate distinct mechanical engagement with the substrate during osmotic challenge. Therefore, the divergent traction force recovery kinetics under hypotonic shock reflect a fundamental difference in how cancer and normal cells mechanically adapt to osmotic stress. Collectively, these results demonstrate that, under osmotic shock, the mechanical behaviors of cancer cells and normal cells are each consistent with their respective intrinsic mechanical responses. Furthermore, they reveal that, beyond these cell-autonomous differences, cancer cells exhibit fundamentally distinct patterns of force interaction with the extracellular matrix compared to normal cells.

**Fig. 2.**
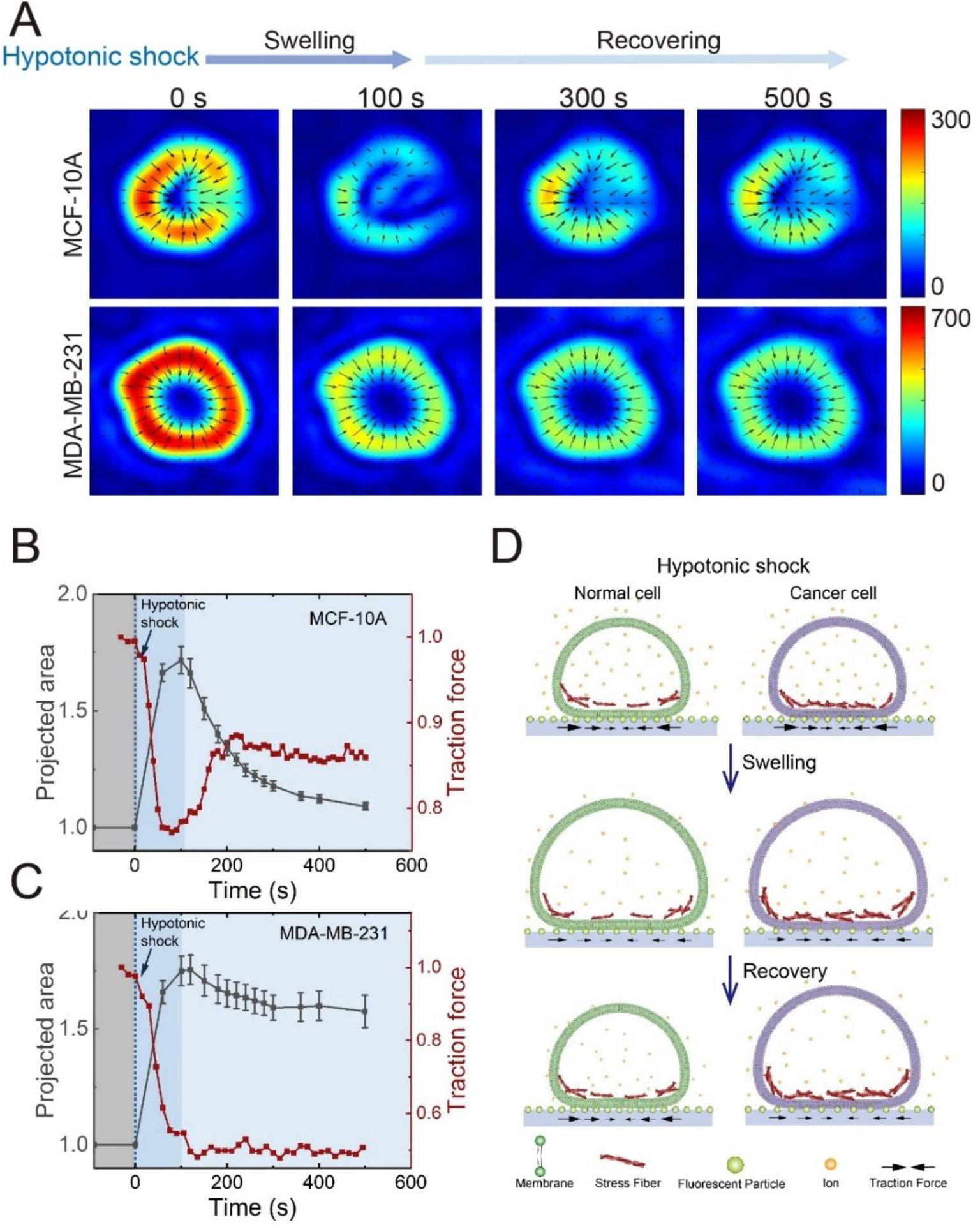
Traction force changes of MCF-10A and MDA-MB-231 under hypotonic shock. (**A**) Representative traction force heatmaps for MCF-10A and MDA-MB-231 cells before and after hypotonic shock (100 mOsm). (**B**) and (**C**) Time course of normalized mean traction force (red line, mean; MCF-10A, n=6; MDA-MB-231, n=7) and projected area (dark grey line, mean ± SE; MCF-10A, n=14; MDA-MB-231, n=15) for (B) MCF-10A and (C) MDA-MB-231 cells following hypotonic shock. (**D**) Schematic illustrating the coupled traction force and volume dynamics in normal versus cancer cells under osmotic shock.

### Actin-mediated Tension as a Determinant of Volume Recovery

Compared to normal cells, cancer cells are softer[20], enabling their stronger invasiveness. Based on these physiological differences and our volume change data, we hypothesized that underlying structural or mechanical variations might account for the differing response patterns between cancerous and normal cells. We characterized the cortical Young’s modulus (Fig. 3A)[25] and the distribution and abundance of actin filaments of both cells. Confocal fluorescence imaging of phalloidin-stained F-actin (Fig. 3B) and 3D reconstruction revealed a concentrated distribution of F-actin within the spherical cap domain of MCF-10A cells. In contrast, MDA-MB-231 cells exhibited a lower density of F-actin in the corresponding region. This higher F-actin density suggested a stiffer cortex and stronger cortical tension. We quantitatively assessed cortical stiffness in both cell lines using atomic force microscopy (AFM). The indentation depth was maintained below 1 μm to ensure the measured Young’s modulus specifically reflected cortical stiffness (Fig. S6A). The Young’s modulus of MCF-10A cells (1.05 ± 0.11 kPa) was higher than that of MDA-MB-231 cells (0.46 ± 0.05 kPa) (Fig. 3D), consistent with the denser F-actin organization observed in MCF-10A cells. Actin structures are key determinants of cellular mechanical behavior. Given this correlation, we hypothesized that the actin-mediated cortical tension might influence the kinetics of cellular volume regulation under osmotic shock. To test our hypothesis, we treated MCF-10A cells with cytochalasin D to inhibit actin polymerization[26], which is expected to reduce actin-mediated tension (Fig. 3C). The cortical Young’s modulus of cytochalasin D-treated MCF-10A cells was notably reduced, and their volume recovery slowed (Fig. 3D and E). Our results show that a reduction in cortical stiffness in MCF-10A cells decreases cortex-mediated tension, leading to slower volume recovery. This provides a cortex-based mechanical explanation for the slower recovery observed in MDA-MB-231 cells, which exhibit lower cortical stiffness than MCF-10A cells. Conversely, MDA-MB-231 cells were treated with jasplakinolide to stabilize actin filaments (Fig. 3C)[27]. We can therefore consider that the cortical tension was enhanced after drug treatment. Although cortical stiffness showed minimal change (Fig. 3D), jasplakinolide-treated MDA-MB-231 cells displayed an accelerated volume recovery rate (Fig. 3E). However, neither drug treatment induced notable changes in the response dynamics to hypertonic shock induced with mannitol solution with the extent of the recovery remaining slight (Fig. 3F). In general, under pharmacological modulation of actin, the dynamics of cellular area response exhibited significant alterations during hypotonic shock, while no such effect was observed under hypertonic shock. Based on this observation, we hypothesized that actin, as a key structural component supporting cellular integrity and generating cortical tension, may follow the same mechanism but the opposite regulating directions through the responding process upon hypotonic and hypertonic shock. This regulatory mechanism of cortical actin is closely linked to cortical tension (Fig. 3G). Under hypotonic shock, cortical tension increases; greater cortical stiffness under these conditions leads to higher tension, which in turn drives faster subsequent volume recovery. In contrast, under hypertonic shock, cells shrink and cortical tension drops sharply. In the absence of active forces that expand the cell, the cell maintains its shrunken morphology, and cortical stiffness has no effect on this process. Furthermore, we measured the cortical stiffness of HepG2 and THLE-2 cell lines. A similar stiffness trend was observed in liver cell lines (Fig. S6B): the cancer cell line HepG2 (0.61 ± 0.04 kPa) displayed lower cortical stiffness than the normal cell line THLE-2 (1.1 ± 0.09 kPa). These results demonstrate that cortical actin content, and thus cortical tension, should play a key role in determining the rate of volume recovery under hypotonic shock. Cells with weaker cortical tension exhibit a slower volume recovery rate, whereas cells with stronger tension recover more rapidly. This mechanical distinction underlies the differential responses observed between cancer and normal cells.

**Fig. 3.**
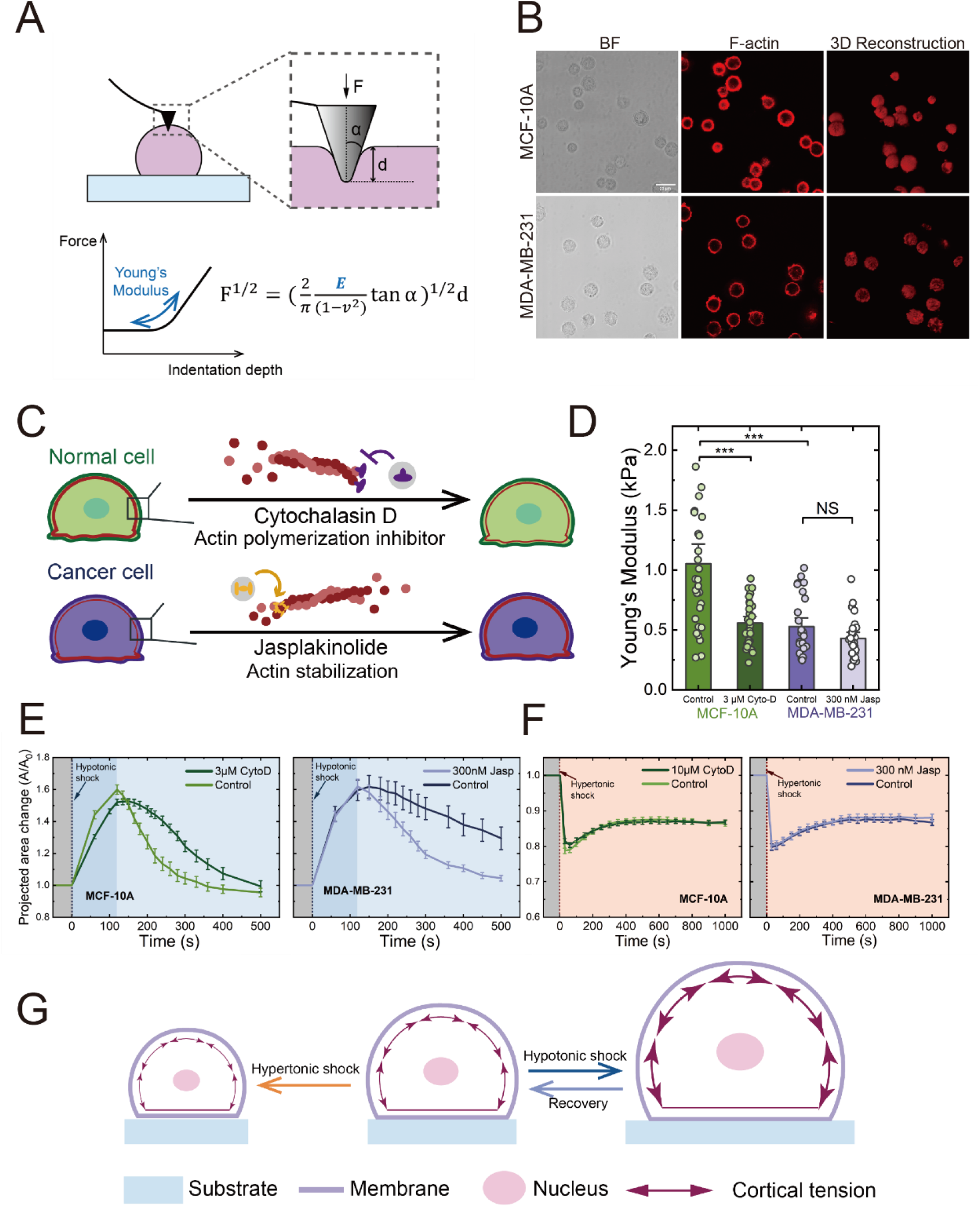
Actin characterization of cancer cells and normal cells. (**A**) Schematic of AFM-based cortical stiffness measurement. (**B**) Representative images of F-actin (phalloidin far-red) with corresponding brightfield, 2D fluorescence, and 3D reconstructed views (Scale bar: 20 μm). (**C**) Schematic of drug mechanisms: jasplakinolide stabilizes actin filaments, cytochalasin D inhibits polymerization. (**D**) Cortical Young’s modulus of untreated and drug-treated cells (untreated controls: MCF-10A, n = 30; MDA-MB-231, n = 26. Drug treated cells: cytochalasin D treated MCF-10A, n = 27; jasplakinolide treated MDA-MB-231, n = 26). (**E**) Projected area change of untreated MCF-10A cells and 3 μM cytochalasin D (untreated controls, n = 14; cytochalasin D treated, n = 11). Projected area change curve of untreated MDA-MB-231 cells and 300 nM jasplakinolide (untreated controls, n = 11; jasplakinolide treated, n = 18). (**F**) Projected area change curve of cells under hypertonic shock mediated by mannitol solution (untreated MCF-10A, n = 28; 10 μM cytochalasin D treated MCF-10A, n = 18; untreated MDA-MB-231, n = 29; 300 nM jasplakinolide treated MDA-MB-231, n = 18). (**G**) Schematic depicting cortical tension changes upon osmotic shock.

### Osmotic Deformation Extent Tuned by Myosin-mediated Contractility

Active cellular contractility is critical for the maintenance of cell shape. Given that actomyosin networks serve as the major components of cellular active contractile machinery[28], we hypothesized that myosin, in addition to actin, also play a role in regulating cellular responses to osmotic shock. We first characterized myosin II in MDA-MB-231 and MCF-10A cells via immunofluorescence staining (Fig. 4A). No significant difference was observed in mean fluorescence intensity between the two cell lines (Fig. 4B). The evidence suggests that myosin did not contribute to the observed differences in osmotic shock response between normal and cancerous cells. To further assess the functional contribution of myosin-driven contractility, we pharmacologically modulated its activity in normal cells. MCF-10A cells were treated with Y-27632, a ROCK inhibitor, to suppress myosin-mediated contractility (Fig. 4C)[29]. Compared with untreated cells, Y-27632-treated MCF-10A cells exhibited a higher maximum projected area upon hypotonic shock (Fig. 4D), indicating that suppressed contractility impaired the cell’s ability to retract, leading to greater peak volume. Conversely, we treated MDA-MB-231 cells with calyculin A to enhance contractility by activating myosin [30]. Calyculin A-treated MDA-MB-231 cells showed that the maximum swelling amplitude was reduced to approximately 1.3 times the initial area, compared to 1.6 times in untreated cells (Fig. 4D). When upon hypertonic shock, Y-27632-treated MCF-10A cells reached a higher minimum projected area than untreated cells. In contrast, calyculin A-treated MDA-MB-231 cells exhibited a significant decrease in the minimum projected area compared to untreated cells. No significant recovery was observed in the drug-treated group or untreated controls. The effects of modulating myosin activity on volume regulation are consistent with its established role in maintaining cellular morphology. The cortex is structured as a network of actin filaments pulled by myosin motors. When myosin activity is inhibited or enhanced, its ability to pull actin filaments changes accordingly. Our results demonstrate that under hypotonic shock, the direction of active contractile forces opposes that of cell swelling, serving to retract the cell and prevent unrestricted expansion. Accordingly, when active contractility is pharmacologically inhibited or enhanced, the peak volume of cell swelling increases or decreases correspondingly. In the case of hypertonic shock, a contractile process is inherently triggered. Therefore, changes in myosin activity directly alter the capacity to pull actin fibers, thereby modulating the extent of volume shrinkage and leading to a higher or lower minimum volume.

**Fig. 4.**
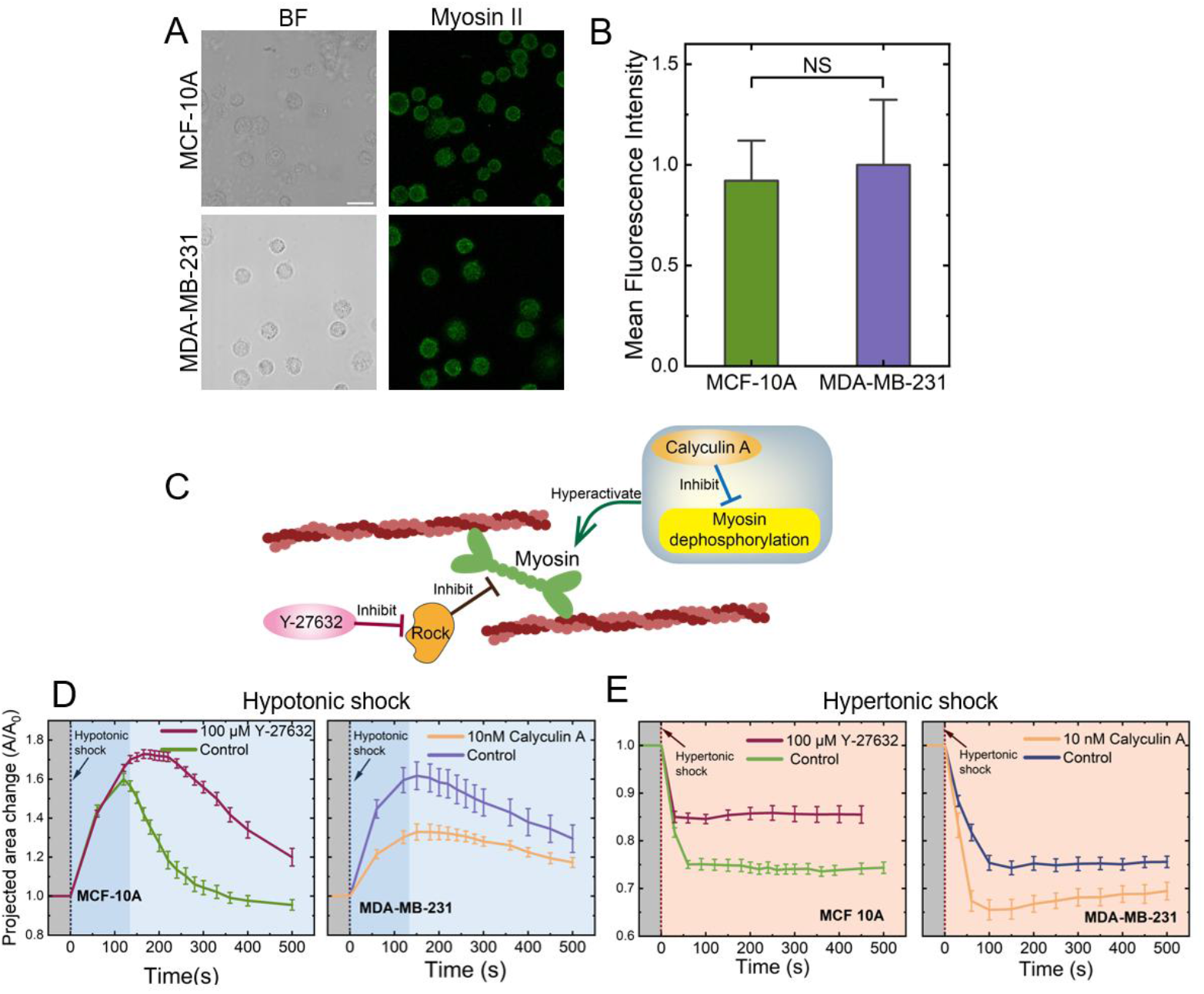
Myosin II characterization and myosin drug perturbation on cells. (**A**) Representative immunofluorescence images of myosin II with corresponding brightfield views (Scale bar: 20 μm). (**B**) Quantification of mean fluorescence intensity for myosin II (MCF 10A, n = 27; MDA-MB-231, n = 23; mean ± SE). (**C**) Schematic of drug mechanisms: Y-27632 inhibits ROCK to reduce contractility, calyculin A activates myosin. (**D**) Projected area change of drug-treated MCF-10A and MDA-MB-231 cells compared to untreated cells (untreated MCF-10A, n = 14; 100 μM Y-27632 treated MCF-10A, n = 17; untreated MDA-MB-231, n = 11; 10 nM calyculin A treated MDA-MB-231, n = 16). (**E**) Projected area change of cells under osmotic shock induced by high concentration PBS solution (untreated MCF-10A, n = 10; 100 μM Y-27632 treated MCF-10A, n = 14; untreated MDA-MB-231, n = 9; 10 nM calyculin A treated MDA-MB-231, n = 12).

### Dynamic Remodeling of Actin under Osmotic Shock

Cancer and normal cells exhibited different recovery rates following hypotonic shock, whereas minimal difference was observed under hypertonic shock. In the process of cellular volume regulation, the membrane tension varies[31]. Cell cortex is a structure that is closely linked to the membrane, as the membrane tension changes, cortical actin senses these alterations and undergoes remodeling[32]. We visualized actin dynamics by labeling actin in both cell types and performing live-cell imaging to track cortical reorganization in real time. Under hypotonic shock, MDA-MB-231 and MCF-10A cells exhibited a similar pattern of actin fluorescence intensity (Fig. 5A and S10): the fluorescence intensity decreased during volume expansion and subsequently increased throughout the volume recovery phase. The observed changes in actin fluorescence intensity cannot be attributed solely to dilution effects resulting from volume changes, as evidenced by distinct morphological alterations in cortical actin staining at the outermost cellular region. In contrast to the inner cytoplasmic area, where fluorescence intensity scaled approximately with volume, the mean intensity of the cortical region did not vary proportionally upon cell swelling or shrinkage. Notably, this outermost layer exhibited a greater extent of intensity fluctuation and underwent concomitant morphological reorganization. This process could be described as actin depolymerization during the cell volume expansion phase, followed by repolymerization during the volume recovery phase. Although the overall dynamic trends were similar, the fluorescence recovery rate was faster in MCF-10A cells than in MDA-MB-231 cells; that is, actin repolymerized more rapidly during recovery in MCF-10A cells (Fig. S10B). The accelerated repolymerization of actin enhanced cortical tension during the recovery phase, thereby expediting volume restoration. This finding further supports our conclusion that actin-driven cortical tension regulates the rate of cell volume recovery. Similarly, in jasplakinolide-treated MDA-MB-231 cells—which showed faster volume recovery—we observed a correspondingly faster rate of actin repolymerization compared to untreated controls after hypotonic shock (Fig. 5A and S10A). This helped explain the faster volume recovery in jasplakinolide-treated cells despite the fact that their cortical stiffness did not increase after drug treatment. Conversely, upon hypertonic shock, actin structures became more condensed, with fluorescence intensity increasing over time as volume decreased (Fig. 5A and S11). This suggests that actin forms a densely packed network to counteract the abrupt osmotic pressure gradient. Subsequently, the cell volume showed minimal recovery, and the decline in fluorescence intensity remained minimal. The distinct behaviors of actin fluorescence intensity aligned with the differential volumetric responses to hypertonic and hypotonic shocks (Fig. 5B). Actin accumulation and repolymerization at the cell cortex facilitates the increase of cellular tension to retract the cell. Our results indicate that cortical actin regulates the volume response to osmotic shock through both static and dynamic mechanisms. Statically, a higher baseline abundance of cortical actin prior to hypotonic shock enhances cortical stiffness, thereby increasing cortical tension and accelerating volume recovery. Dynamically, accelerated actin repolymerization facilitates a more rapid buildup of cortical tension during the recovery phase. These two pathways exert additive effects in the same direction to modulate the rate of cell volume recovery. Conversely, under hypertonic shock, cortical tension decreases as the cell shrinks, and cortical actin reorganizes into a dense layer. In the absence of a sufficient driving force to expand the cell back to its original volume and morphology, neither the baseline abundance of cortical actin nor the rate of actin polymerization exerts a significant effect on the cellular response. Together, our experimental findings demonstrate that the regulation of cortical mechanical behavior is critical for cellular adaptation to osmotic shock.

**Fig. 5.**
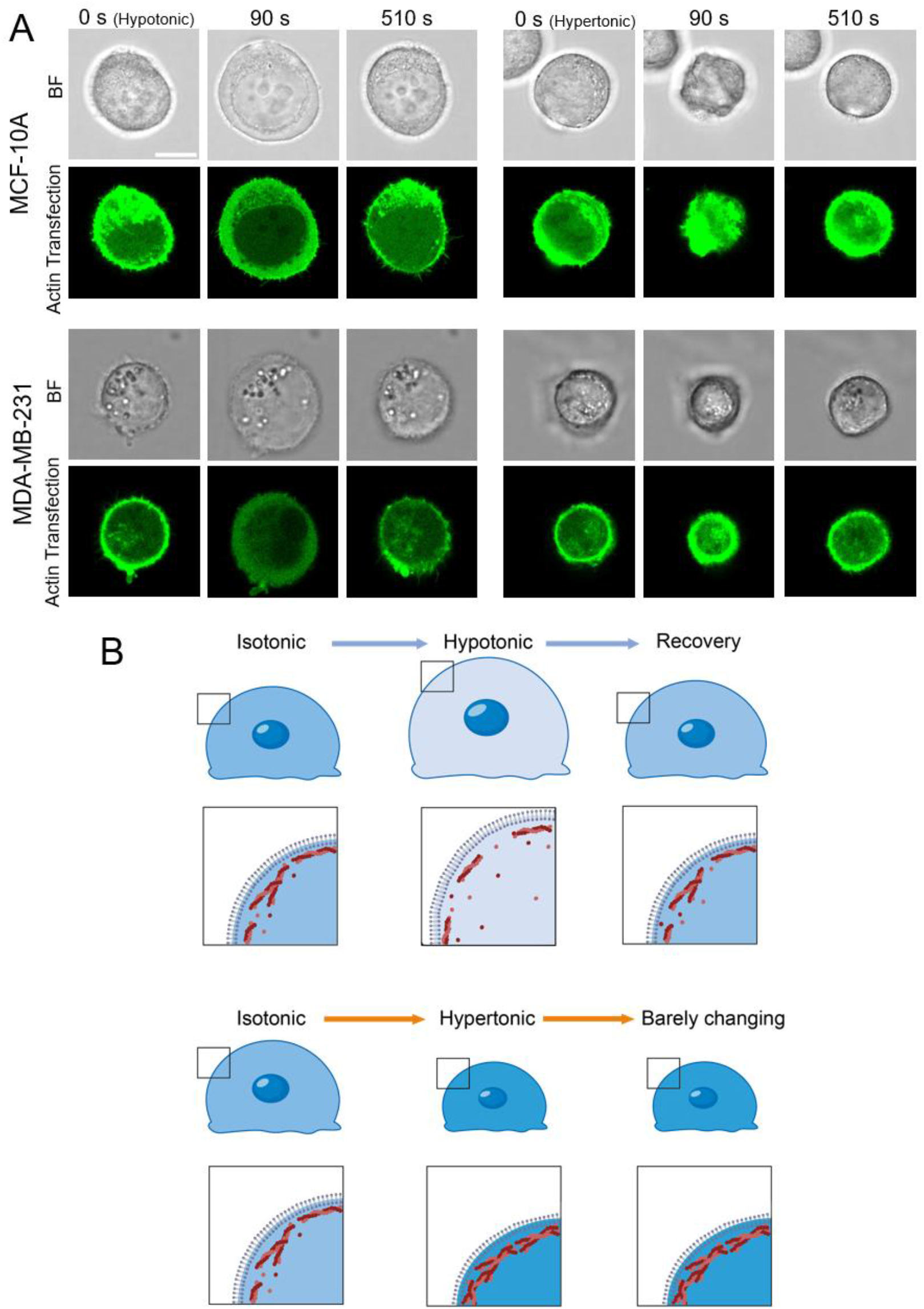
Myosin II characterization and myosin drug perturbation on cells. (**A**) Visualization of F-actin remodeling dynamics in MCF-10A and MDA-MB-231 cells under hypotonic and hypertonic shock (Scale bar: 10 μm). (**B**) Schematic summarizing actin network states under iso-, hypo-, and hypertonic conditions.

### Effects of Substrate Stiffness on Recovery Speed upon Hypotonic Shock

Under slightly adherent conditions, cells attach to the substrate, and cellular deformation exerts mechanical forces on the substrate. Conversely, the substrate exerts a reaction force on the cell; therefore, its mechanical properties are likely to influence cellular behavior. To explore the influence of substrate mechanical properties on osmotic responses, we prepared hydrogels with stiffnesses of 3, 23, and 70 kPa and exposed slightly adherent C2C12 cells to hypotonic shock. We found that the stiffness of the substrate significantly influenced how slightly adherent cells responded to osmotic shocks. As substrate stiffness increased, the rate of volume recovery decreased (Fig. 6A). A similar trend was observed in slightly adherent MDA-MB-231 cells cultured on hydrogels with stiffnesses of 3 kPa and 70 kPa (Fig. S12). We also exposed MDA-MB-231 cells on substrates of 3, 23, and 70 kPa to hypertonic shock induced by high-concentration PBS (Fig. 6B). No notable differences were detected in the minimum volume or the recovery dynamics across substrate stiffnesses. To gain insight into the variation in volume recovery rate under hypotonic shock, we stained actin filaments in C2C12 cells on substrates of 3, 23, and 70 kPa (Fig. 6C). The results showed that, given the same adhesion time, C2C12 cells formed a greater number of stress fibers in the basal contact region on stiffer substrates than on softer ones. A similar phenomenon was observed for MDA-MB-231 and MCF-10A cells. MDA-MB-231 cells formed a greater number of stress fibers in the cell-substrate contact region compared to MCF-10A cells. A denser network of stress fibers on the substrate may impose a stronger constraint on cellular deformation during the response to hypotonic shock, thereby slowing the subsequent volume recovery process. The effect of substrate stiffness on osmotic response demonstrates that ECM mechanical properties profoundly influence cellular behavior upon hypotonic shock; whereas upon hypertonic shock, there exists no notable difference, which is consistent with all the former results. Collectively, these observations support the concept of the opposite directions of cellular responding system to hypertonic versus hypotonic shock and offer a more complete explanation for the differential volume recovery rates under hypotonic conditions.

**Fig. 6.**
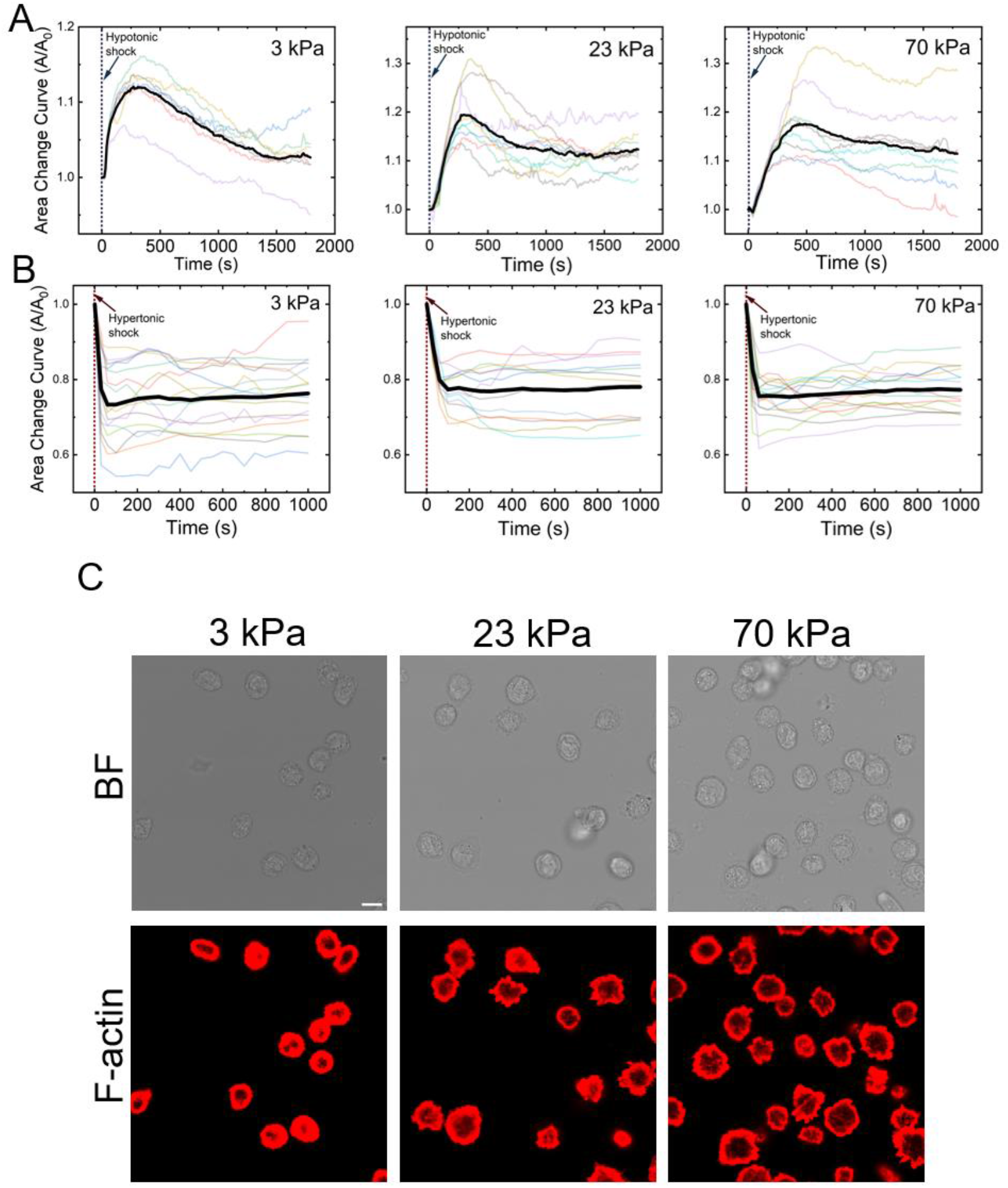
Substrate stiffness modulates osmotic response. (**A**) Normalized projected area dynamics of C2C12 cells upon hypotonic shock on substrates with different stiffness (3 kPa, n = 6; 23 kPa, n = 9; 70 kPa, n = 8). (**B**) Normalized projected area dynamics of MDA-MB-231 cells upon hypertonic shocks on substrates with different stiffness (3 kPa, n = 13; 23 kPa, n = 16; 70 kPa, n = 17). (**C**) Representative basal view of phalloidin-stained F-actin in slightly adherent C2C12 cells cultured on hydrogels with Young’s moduli of 3, 23, and 70 kPa (Scale bar: 10 μm).

### Theoretical Modeling of Cellular Response to Osmotic Shock

To unify the factors underlying the osmotic response variations observed in our experiments, we developed a theoretical model incorporating cell shape regulation, cortical tension[33], ion transport[16], cellular adhesion[34], and energy conversion (Fig. 7A). In this model, the cytoplasm is surrounded by the cell membrane and the cell cortex. The cell cortex is a layer of actomyosin network attached to the inner side of the cell membrane. Given that the cortex and membrane remain coupled within a certain range of osmotic pressure, these two structures are simplified as a single composite shell in the model[16]. The physical parameters of this composite shell, such as Young’s modulus, are therefore approximately equivalent to those of the cell cortex. The cell membrane is embedded with ion channels for active transport and passive transport, which collectively regulate the dynamic response of ion concentrations under osmotic shock. The permeability of active transport ion channels is affected by factors such as the transmembrane osmotic pressure difference and ATP concentration. The permeability of passive transport channels, meanwhile, is primarily regulated by cortical tension. Transmembrane water flux is modulated by both the osmotic pressure difference and the hydrostatic pressure. In adherent cells, focal adhesions connect the cell to the substrate through ligand-receptor binding, transmitting cellular contraction forces and mediating force variations induced by osmotic shock.

**Fig. 7.**
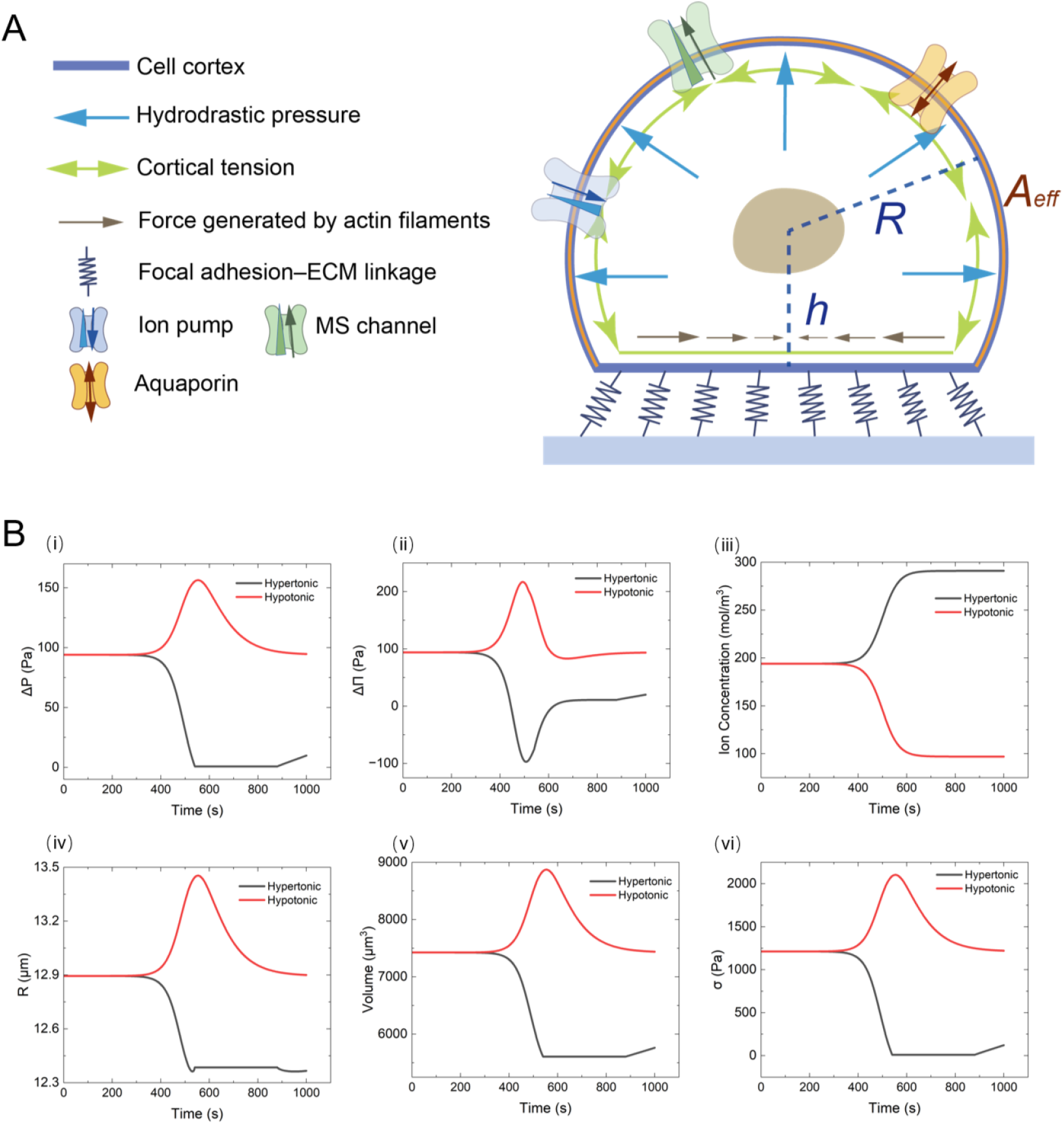
Theoretical model and simulation outputs. (**A**) Schematic of the simplified model for a slightly adherent cell responding to osmotic shock. Key components include the composite membrane-cortex shell, ion and water transporters, and dynamic adhesion sites. (**B**) Simulated temporal changes in key parameters under hypertonic (black) and hypotonic (red) shocks: (i) transmembrane hydrostatic pressure difference (*ΔP*), (ii) transmembrane osmotic pressure difference (*ΔΠ*), (iii) intracellular ion concentration, (iv) cell radius, (v) cell volume, and (vi) cortical stress.

### Dynamics of Adhesion Energy

Distinct from suspended cells, adherent cells exhibit morphological variations that arise from the combined effects of volumetric alterations and shape changes. Our model focuses on simulating the dynamic adhesion energy during cell attachment and detachment, integrating cellular mechanics and transport phenomena (Fig. 7A).

The adhesion energy density *Γ* is not constant during the dynamic process of cell-substrate interaction[35, 36]. As the cell is pulled away from the substrate, the adhesion contact radius *r*_*c*_(*t*) decreases, signifying a weakening of the adhesive bond. To simulate this, we proposed a phenomenological model based on the physical picture of a “pre-contracted” adhesive patch existing within an energy potential well. We defined two key geometric states for the contact radius, the hypothetical rest state and the stable equilibrium state.

The hypothetical rest state, where the cell contacts the substrate with zero adhesion energy, represents the boundary of the stable adhesion zone. The stable equilibrium state, where the adhesion energy density is at its maximum *Γ*_0_, is the actual mechanically stable contact state which is also the cell’s initial state. The transition from the larger rest radius to the smaller stable radius is viewed as a “pre-contraction”[37]. We quantified this with a dimensionless parameter *ϵ*:

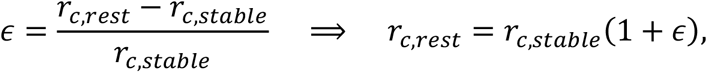

where *r*_*c,rest*_ denotes the contact radius under the hypothetical rest state and *r*_*c,stable*_ denotes that under the stable equilibrium state. Adhesion energy is represented using a quadratic potential, with the assumption that the stored energy is proportional to the square of the deviation from the zero-energy rest state:

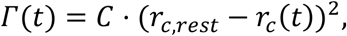

where *C* is a constant and *r*_*c*_(*t*) is the contact radius at any given time. We determined the constant *C* by applying the condition that at the stable state, *Γ*(*t*) = *Γ*_0_ when *r*_*c*_(*t*) = *r*_*c,stable*_:

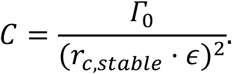

Substituting the expressions for *C* and *r*_*c,rest*_ back into the energy model yields the final equation used in our simulation:

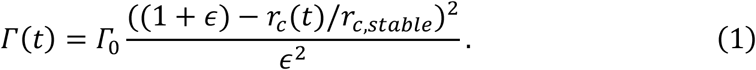

### Molecular Basis: Adhesion Kinetics

The maximum adhesion energy density *Γ*_0_ and the stability of the bond are grounded in molecular kinetics. Adherent cells interact with the substrate via focal adhesions, where integrins form ligand-receptor complexes[36].

Cell adhesion to the extracellular matrix is a fundamental biological process whose thermodynamics can be described by classical physicochemical principles. When cells undergo slight adhesion to a substrate surface, the adhesion energy density *Γ* at the resulting focal adhesions can be approximated by the Young-Dupré equation[34]:

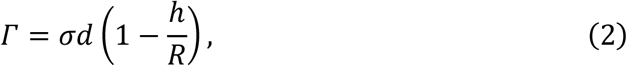

where *d* is the thickness of the cell cortex, *σ* is the cortical stress, *R* represents the radius of the cell, and *h* denotes the distance from the spherical center of the cellular dome to the adhesion plane. The molecular bonds mediating cell-substrate adhesion can be modeled as a system of distributed discrete springs. The adhesion energy density *Γ* relates to the density of engaged molecular bonds *ρ* and their extension *l*:

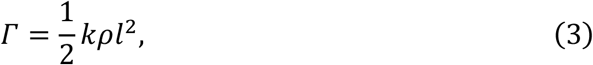

where the density of engaged bonds, *ρ*, is determined by the dynamic turnover of ligand-receptor binding (see Supplementary Note A for kinetic equations). To capture the non-trivial response of bond lifetime to force, we employed a two-pathway model describing catch-slip bond behavior[38]:

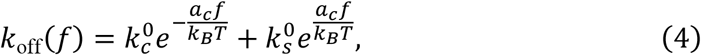

in which 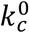 and 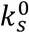 are the zero-force dissociation rates for the catch and slip pathways, respectively, and *a*_*c*_ are their characteristic reactive distances, while *k*_*B*_ and *T* represent the Boltzmann constant and absolute temperature. This model posits that the bond can dissociate via two competing pathways: a “catch bond” pathway stabilized by force, and a “slip bond” pathway destabilized by force.

### Coupling with Morphological Regulation

The dynamic adhesion energy defined in Eq. (1) relies heavily on the contact radius *r*_*c*_(*t*), which evolves as the cell undergoes volume regulation and morphological changes. Therefore, simulating cellular morphological changes is central to our model. We developed an integrated framework incorporating water transport, cortical tension dynamics, and ion transport to simulate cell morphology dynamics.

1. Water Transport and Volume Change The cell volume *V* changes due to transmembrane water transport driven by hydrostatic (*ΔP*) and osmotic (*ΔΠ*) pressure differences[39]:

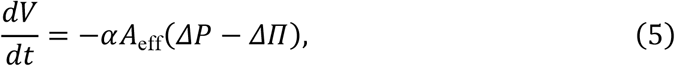

where α is the water transport rate constant characterizing membrane permeability to water molecules, and *A*_eff_ is the effective surface area of the cell excluding the adherent region. *ΔP* = *P*_in_ − *P*_out_ and *ΔΠ* = *Π*_in_ − *Π*_out_ respectively represent transmembrane hydrostatic pressure and osmotic pressure differences. As the cell swells or shrinks, the effective surface area *A*_eff_ and the contact geometry (*R* and *h*) should shift, altering *r*_*c*_(*t*) and thus the adhesion energy *Γ*(*t*). (See Supplementary Note B for geometric derivations).
2. Cortical Stress The hydrostatic pressure is balanced by the cortical stress *σ*. The cell cortex behaves as an elastic material subject to both passive deformation stress *σ*_*passive*_ and active myosin contractility *σ*_*a*_[33]:

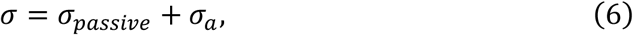

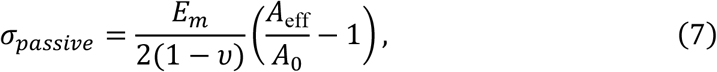

where *E*_*m*_ represents the elastic modulus of the cell cortex, *A*_eff_ and *A*_0_ denote the effective surface area of the cell excluding the adherent region and its natural (stress-free) area respectively, and υ is the Poisson’s ratio.
3. Ion Transport Regulation

The osmotic pressure difference *ΔΠ* is regulated by active and passive ion transport. Mechanosensitive (MS) channels open in response to membrane stress, allowing passive ion flux to relieve osmotic imbalances[16]. (See Supplementary Note C for transport equations).

### Theoretical Simulation Results

Solving the coupled equations, we found that our simulation results matched the experimental results presented earlier. The extracellular osmotic pressure changes (*Π*_out_) were set as the stimulus in the numerical theoretical model (Fig. S13A). Upon hypotonic shock, the simulated cell exhibited a reduction in intracellular osmotic pressure, and the transmembrane osmotic pressure difference showed an initial increase followed by a decrease (Fig. 7B (ii)). To compensate for the initial increase in osmotic pressure difference, the simulated cells transported ions outward (Fig. 7B (iii)). Our simulation results matched this description. Furthermore, the transmembrane hydrostatic pressure difference also showed an initial rise followed by a decline (Fig. 7B (i)). This pattern arose because the cells first absorbed water and swelled, then lost water and contracted. The simulated changes in cell radius, volume, and cortical stress aligned with our experimental observations (Fig. 7B (iv)-(vi)). Under hypertonic shock, the parameter changes followed a process that also closely matched the experiments. Cells underwent volume reduction immediately after hypertonic shock, expelling water and transporting ions inward (Fig. 7B (i)-(iii)). The simulated minimal volume recovery under hypertonic shock is consistent with our experimental hypothesis that this phase involves a physical opposition to actomyosin-driven contraction. The simulation data show a sharp decline in cortical stress and hydrostatic pressure difference upon hypertonic shock with minimal recovery, indicating that there would be not enough force for cell to regain the volume. This provides supporting evidence for our experimental hypothesis.

Our experimental results demonstrated that normal cells (MCF-10A and THLE-2) had a higher Young’s modulus than cancer cells (MDA-MB-231 and HepG2), and that this stiffness difference reduced cellular tension and consequently slowed volume recovery under hypotonic shock. Simulations showed that, with other parameters fixed, the recovery rate increased with cortical stiffness (Fig. 8A). Specifically, the time required to recover to 60% of the peak expanded area progressively decreased as the cortical elastic modulus increased (Fig. 8B). Simulations under hypertonic shock also matched the experiments: the extent and rate of recovery changed minimally with increasing cortical stiffness (Fig. 8E). A consistent pattern was observed in which cortical stress dropped sharply and showed negligible recovery, regardless of cortical Young’s modulus. This universal stress response provides a mechanistic explanation for the lack of significant volume recovery across varying cortical stiffness upon hypertonic shock. To further explore the mechanism underlying the differential recovery, we calculated changes in hydrostatic pressure difference and cortical stress for different values of cortical Young’s modulus (Fig. 8C and D). The hydrostatic pressure difference drives water efflux from the intracellular to the extracellular compartment, while cortical stress actively expels water from the cell. Thus, during the recovery phase, cortical stress served as the key driver of outward water flow. Greater cortical stress enhances the cell’s capacity for water expulsion, thereby accelerating the efflux rate and promoting faster volume recovery. Simulations indicated that cells with a stiffer cortex exhibited a more pronounced increase in hydrostatic pressure upon swelling (Fig. 8C), leading to accelerated water efflux and faster volume restoration. Correspondingly, these cells also reached a higher peak cortical stress (Fig. 8D), which further augmented the driving force for water expulsion, collectively accelerating recovery.

**Fig. 8.**
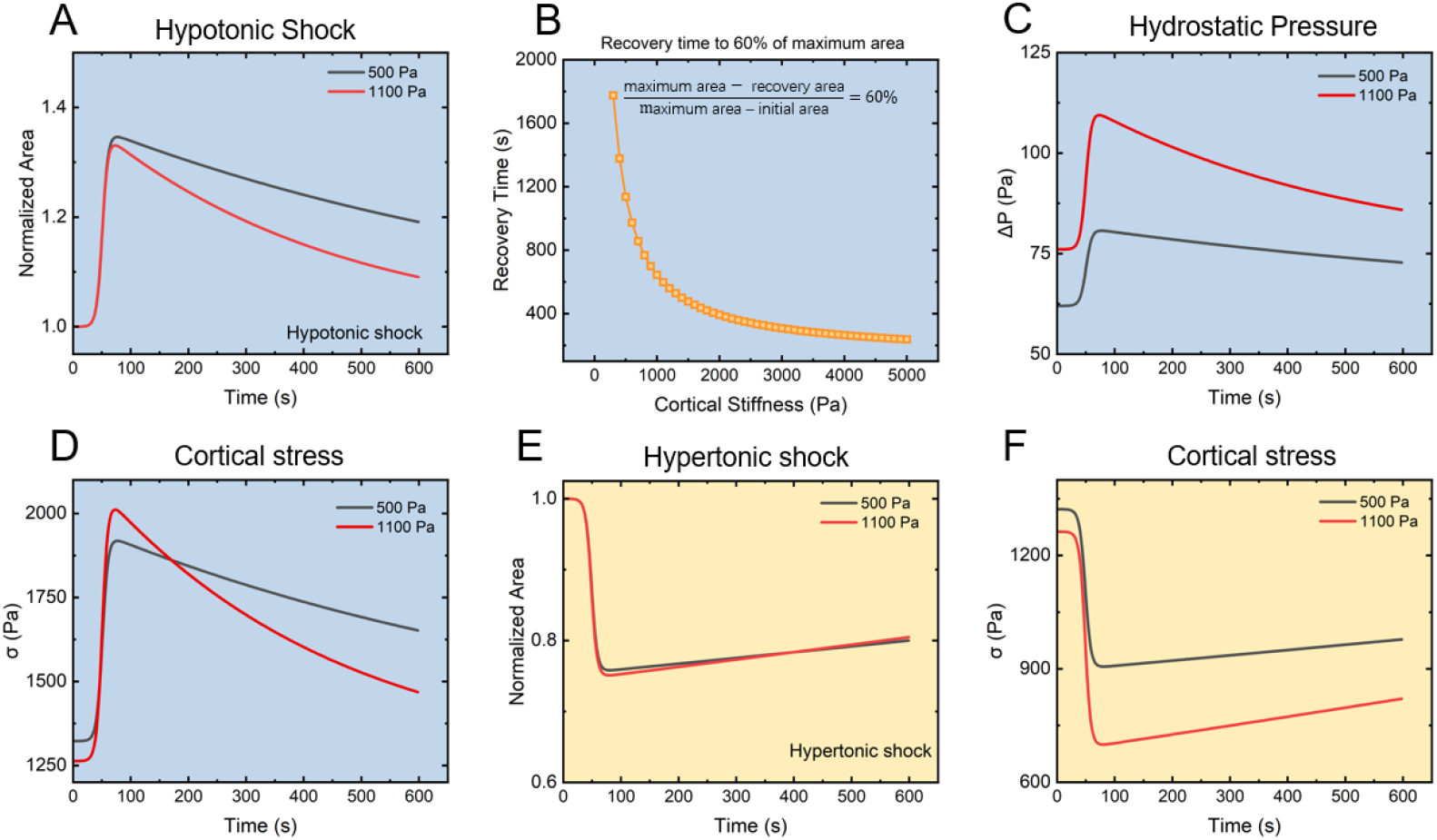
Model predictions on the role of cell cortex (dark gray line: 500 Pa; red line: 1100 Pa). (**A**) Schematic of the simplified model for a slightly adherent cell responding to osmotic shock. Key components include the composite membrane-cortex shell, ion and water transporters, and dynamic adhesion sites. (**B**) Simulated temporal changes in key parameters under hypertonic (black) and hypotonic (red) shocks: (i) transmembrane hydrostatic pressure difference (*ΔP*), (ii) transmembrane osmotic pressure difference (*ΔΠ*), (iii) intracellular ion concentration, (iv) cell radius, (v) cell volume, and (vi) cortical stress.

Overall, our simulation results are in good agreement with the experimental results. The model we established provides theoretical support for the experimental conclusions, confirming that the cortical mechanical property modulates the cellular response rate to osmotic shocks.

## Discussion

Our study elucidates the differences in volume recovery between normal and cancer cells following hypotonic shock and delineates the underlying regulatory system that governs this divergent osmotic response. We focused on two coupled readouts, cellular volume changes and traction force changes. The dynamics of these two parameters were closely coupled. While both normal (MCF-10A, THLE-2) and cancer (MDA-MB-231, HepG2) cells initially swell to a similar extent, their recovery trajectories sharply bifurcate: normal cells rapidly regain original volume and traction force, whereas cancer cells recover slowly with traction force remaining slightly changed. This disparity is rooted in the mechanical capacity of the submembrane actomyosin cortex. Quantitative characterization shows that the stiffer, denser cortex of normal cells generates greater actin-mediated tension, driving efficient regulatory volume decrease (RVD). Conversely, the softer cortex of cancer cells generates weaker cortical tension, resulting in sluggish recovery. Pharmacological modulation solidified this causality—disrupting actin in normal cells slowed recovery, while stabilizing actin in cancer cells accelerated it—confirming that actin-driven tension is a key determinant of recovery kinetics. Real-time imaging of F-actin dynamics reinforced this link, showing that depolymerization during swelling and repolymerization during recovery were synchronized with volume changes and volume recovery was faster when this repolymerization process occurred in a more rapid way. Beyond actin structures on the cortex, myosin-mediated force dynamically tunes the response amplitude. As the myosin levels were similar in cancer cells and normal cells, their swelling amplitudes were on the same level as well. When myosin-mediated contractility was modulated, the swelling amplitude changed: inhibition increased the maximum volume, whereas enhancement reduced it (Fig. 9).

**Fig. 9.**
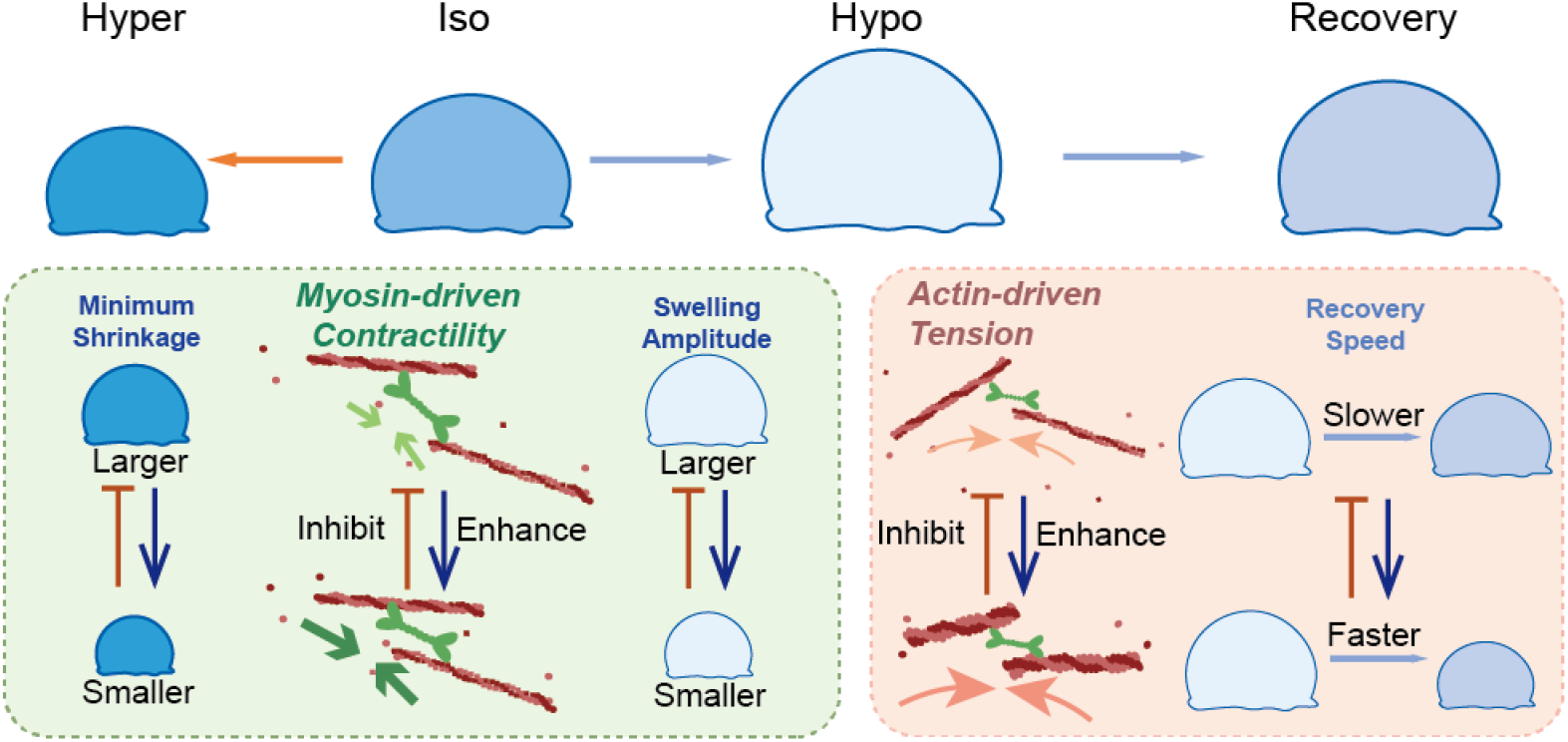
Diagram depicting the mechanism by which the cell cortex modulates volume adaptation to osmotic stress. More precisely, the speed of cellular contraction is controlled by actin-based cortical tension, whereas the magnitude of contraction is regulated by myosin-generated active forces.

A striking asymmetry emerged between responses to hypotonic and hypertonic stress. While hypotonic recovery was actively driven by the actomyosin cortex, hypertonic shrinkage and any subsequent minimal recovery were indistinguishable between cell types. These distinct patterns indicate that the cortex may respond in opposite directions to hypo- and hypertonic shock. In contrast to hypotonic shock—which increases cortical stress and triggers transient actin depolymerization followed by reassembly—hypertonic shock leads to dissipation of cortical stress and concurrent actin accumulation, forming a denser, more static architecture. Within this structure, myosin generates contractile forces by pulling on actin filaments. This explains why pharmacological perturbations of actin or myosin specifically altered hypotonic responses while leaving hypertonic ones unchanged.

The regulatory system extends beyond cell-intrinsic properties to integrate signals from the mechanical microenvironment. Substrate stiffness emerged as an external modulator, where stiffer substrates, promoting robust stress fiber formation, slowed hypotonic recovery by enhancing cellular constraint. This effect was absent under hypertonic conditions, further supporting the concept of opposite directions of cortical roles in responses to hypo-versus hypertonic shock. To unify these principles, we developed a theoretical model coupling water transport, cortical mechanics, and dynamic adhesion. The model successfully captured the experimental observations: stronger cortical tension generates higher hydrostatic pressure and cortical stress during swelling, driving faster water efflux and recovery, and it correctly reproduces the minimal influence of cortical or substrate stiffness on hypertonic responses.

Collectively, our findings establish that the differential osmotic recovery capacity stems from a disparity in the efficacy of the cortex regulatory system. This system integrates intrinsic mechanical properties with extrinsic mechanical cues from the substrate to govern volume homeostasis after osmotic shock. The significantly slower and incomplete volume recovery observed in cancer cells may thus provide a mechanistic explanation for the easier rupture of cancer cells under extreme hypotonic environments documented in prior clinical studies. The compromised cortex in cancer cells might delay volume restitution and create deeper cellular damage. This mechanistic vulnerability could provide an explanation for the therapeutic efficacy of extreme hypotonic conditions in cancer treatment—an effect leveraged in adjuvant peritoneal lavage with distilled water in surgical oncology. Our investigation into the differences in cortical mechanical properties between normal and cancer cells, and how these differences influence cellular responses to osmotic stress, may provide osmotic insights and a cellular mechanics reference for the future design of cancer therapeutic strategies. Furthermore, our findings on how substrate stiffness modulates cellular osmotic responses contribute to understanding the mechanobiological behavior of cells under osmotic stress variation when the mechanical properties of the extracellular matrix change in vivo—such as during disease progression or pharmacological treatment. Thus, our work links a fundamental mechanobiological deficiency to a clinically observed therapeutic window, offering a biophysical rationale for targeting osmotic homeostasis in cancer therapy.

## Materials and Methods

Detailed information on cell culture, osmotic shock experiments, traction force microscopy, pharmacological treatments, atomic force microscopy measurements, and theoretical modeling is provided in Supplementary Information, Materials and Methods. Briefly, projected area changes of MDA-MB-231 and MCF-10A cells were recorded in real time under hypo- and hypertonic shocks. Traction forces were quantified using substrate displacement analysis. Cortical stiffness was measured by AFM with an indentation depth of <1 μm. Actin and myosin perturbations were performed using cytochalasin D, jasplakinolide, Y-27632, and calyculin A. A theoretical model coupling adhesion energy, ion/water transport, and cortical mechanics was developed as described in Supplementary Note A, B and C. Statistical analyses were performed using OriginPro 2024. Data are presented as mean ± SEM. Comparisons between two groups were conducted using two-tailed, two-sample t-test. P<0.05 was considered statistically significant. All experiments were repeated at least three times independently, with details provided in the figure legends and Supplementary Information. All numerical simulations were performed using MATLAB.

## Acknowledgments

This work was supported by the National Natural Science Foundation of China.

National Natural Science Foundation of China grant 12232017 (QZ, SW) National Natural Science Foundation of China grant 12222212 (QZ, SW)

## Author contributions

Conceptualization: QZ, SW

Methodology: XW, JG, WW, SZ, TZ

Investigation: XW, WW, CW, YC, TZ, SZ

Formal Analysis: XW, WW

Writing – Original Draft: XW

Writing – Review & Editing: XW, QZ, SW

Supervision: QZ, SW

Project Administration: QZ, SW

## Competing interests

The authors declare no competing interest.

## Notes

### Competing Interest Statement

The authors have declared no competing interest.

