## Supplementary Text and Figures for "Volume regulation of cancer cells during osmotic pressure variation"

**Materials and Methods**

**Cell Culture.**

MCF-10A cells (Procell) were cultured with MCF-10A-specific medium (Procell). THLE-2 cells (Procell) were cultured with THLE-2-specific medium (Procell). MDA-MB-231 cells were cultured with 1640 high-glucose medium (Gibco) containing 10% fetal bovine serum (FBS, Gibco), penicillin (100 U/mL, Gibco), and streptomycin (100 μg/mL, Gibco). HepG2 cells and C2C12 cells were cultured with Dulbecco’s modified Eagle’s medium (DMEM, Gibco) containing 10% fetal bovine serum (FBS, Gibco), penicillin (100 U/mL, Gibco), and streptomycin (100 μg/mL, Gibco). All cells were cultured in a 37 °C incubator with a 5% CO₂ atmosphere. Cells were detached from culture flasks using 0.25% trypsin (Gibco) containing ethylenediaminetetraacetic acid (EDTA). Before osmotic shocks, cells were resuspended, seeded onto hydrogel substrates, and cultured for 20 minutes at 37 °C with 5% CO₂ to establish slightly adherent conditions.

**Drug Treatment**

Cells were preincubated with the following conditions before resuspension. All drugs were dissolved in the respective culture medium matched to each cell line. The following pharmacological inhibitors were preincubated with the following conditions: 3 μM cytochalasin D for 30 min (MCE), 300 nM jasplakinolide for 1 h (MCE), 100 μM Y-27632 for 30 min (MCE), 10 nM calyculin A for 30 min (Macklin).

**Osmotic Shock**

Cells were allowed to adhere lightly for 20 minutes. The culture medium was then removed and replaced with phosphate-buffered saline (PBS) at 300 mOsm, which approximates the osmotic pressure of human blood plasma. Hypotonic shock was induced by adding distilled water to achieve a final osmolarity of 100 mOsm or 200 mOsm. Hypertonic shock was induced using either 2.5× PBS or a mannitol solution prepared in 1× PBS (944 mOsm), yielding final osmolarities of 450 mOsm or 600 mOsm.

**Fluorescence Staining**

Cells were incubated on hydrogel for 20 min to maintain slightly adherent conditions before being fixed with 4% (w/v) paraformaldehyde in PBS and permeabilized with 0.1% Triton X-100 in PBS. Cells were blocked with 1% bovine serum albumin (BSA)/PBS for 1 hour at room temperature (RT) prior to antibody exposure. Then cells were incubated with anti-paxillin primary antibody (1:50, Cell Signaling) overnight at 4 °C, followed by Alexa Fluor 488-conjugated secondary antibody (1:500, Abcam) for 1 h at RT. MCF-10A and MDA-MB-231 cells were processed similarly, except that the primary antibody was replaced with myosin II antibody.

**Young’s Modulus Measurement**

Young’s modulus data of cells were acquired in PBS solution using the atomic force microscopy Nanowizard Sense+ (Bruker) mounted on a Leica DMi8 inverted epifluorescence microscope. Contact Mode was performed using silicon nitride cantilevers (DNP-D, Brucker) with a nominal spring constant of 0.06 N/m and a pyramidal tip with a nominal half angle of 18.7°. The tip has a radius of 20 nm. In order to obtain reliable data, we calibrated the probe's spring constants using the Sader method. To ensure the measured Young’s modulus reflected the cell cortex rather than the cytoplasm or nucleus, setpoints were relatively small in the range of 0.4-0.6 nN. The indentation depths were therefore controlled within 1 μm. Approach-retract force-indentation curves were recorded and analyzed with the Hertz-Sneddon model to calculate the Young’s modulus.

**Elastic Substrate Fabrication**

The glass-bottom culture dishes were pretreated with 4% (v/v) 3-Aminopropyltrimethoxysilane (APTES) solution before being treated with 0.5% glutaraldehyde solution. Culture dishes were dried in air after being washed with deionized water. Solutions of 5% (w/v) acrylamide, 0.5% (w/v) N,N-methylene-bis-acrylamide, 0.1% (v/v) N,N,N’,N’-tetramethylethylenediamine and 0.05% (w/v) ammonium persulfate were mixed into pre-gel solution. The pre-gel solution was deposited onto pre-cleaned circular quartz plates. The solution between the culture dishes and quartz plates flattened under gravity and polymerized at room temperature for 40 minutes. After quartz plates removal, the polyacrylamide hydrogel was immersed in deionized water to eliminate unreacted monomers, yielding substrate-ready hydrogels. This method produced hydrogels with a Young’s modulus of approximately 10 kPa. To enhance cell adhesion on hydrogel surfaces, the crosslinker sulfo-SANPAH was used to facilitate covalent conjugation between type I collagen and polyacrylamide hydrogels. The dishes were then exposed to UV light (10W) for 30 minutes, followed by overnight coating with 200 μL of 0.6% (v/v) rat tail type I collagen solution (Thermo Fisher, diluted in 6 mM acetic acid) at 4°C. Elastic hydrogels were fabricated using the same procedure, with stiffness variations achieved exclusively through adjusted monomer ratios as specified below: 3 kPa, 5% (w/v) acrylamide, 0.1% (w/v) N,N-methylene-bis-acrylamide, 0.1% (v/v) N,N,N’,N’-tetramethylethylenediamine and 0.1% (w/v) ammonium persulfate; 23 kPa, 10% (w/v) acrylamide, 0.2% (w/v) N,N-methylene-bis-acrylamide, 0.1% (v/v) N,N,N’,N’-tetramethylethylenediamine and 0.1% (w/v) ammonium persulfate; 70 kPa, 10% (w/v) acrylamide, 0.5% (w/v) N,N-methylene-bis-acrylamide, 0.1% (v/v) N,N,N’,N’-tetramethylethylenediamine and 0.1% (w/v) ammonium persulfate.

**Traction Force Microscopy**

To fabricate hydrogels embedded with fluorescent beads for traction force microscopy, the following steps were added prior to sulfo-SANPAH treatment: a mixture of 4 μL green fluorescent particle solution (2% w/v, 0.5 μm, carboxyl-modified) and 200 μL deionized water was added to 10 kPa hydrogel-coated dishes for 40 minutes, followed by three washes with deionized water. Later, the dishes were treated with a 0.5% (w/v) NHS and 2% (w/v) EDC mixture for 2 hours, then carefully rinsed three times with deionized water. The cell adhesion promotion procedure was then repeated on the hydrogel surfaces. Cells were cultured on hydrogels embedded with fluorescent particles for 20 min before the osmotic shocks. Z-stack imaging of surface-conjugated fluorescent particles was performed using confocal microscopy, with subsequent image superposition to generate computational speckle patterns. For cell-seeded samples, initial speckle patterns were recorded as deformation maps. Following cell removal via 0.5% (w/v) sodium dodecyl sulfate (SDS) treatment, the same regions were rescanned to obtain reference speckle patterns. Hydrogel surface displacements were calculated based on images of beads using digital image correlation in the PMLAB software (Nanjing PMLAB Sensor Tech Co.). The displacement data were imported into the open-source software PYTFM for traction force inversion. This software performs force reconstruction based on the Boussinesq equation, with computations conducted in Fourier space. For traction force quantification, the algebraic sum of traction forces at all measurement points was calculated. Then mean traction force on each point was obtained for statistical analysis.

$$F_{Average}=\frac{\sum_{i=1}^{n} \sqrt{F_{x}^{2}\left( i \right)+F_{y}^{2}\left( i \right)}S(i)}{\sum_{i=1}^{n} S(i)}$$

**Fluorescent Labeling of Actin Filaments**

To generate a GFP-actin stable cell line, plasmids of pLVX-GFP-actin were cotransfected with two packaging plasmids, psPAX2 and pMD2.G, in HEK293T cells to produce lentivirus. Two days after transfection, the supernatant containing lentivirus was collected and filtered through a 0.45 μm filter. MCF-10A and MDA-MB-231 cells were infected with the lentivirus in the presence of 8 μg/mL polybrene (Sigma) for 12 h. Twenty-four hours after transfection, puromycin (2 μg/mL) was added to select the successfully transduced cells for another 72 h.

**Supplementary Notes and Figures**

**Note A:** Adhesion Kinetics Detail

The rate of change of bond density $\rho$ is described by:

$$\frac{d\rho}{\mathrm{dt}}=k_{\mathrm{on}}\left( \rho_{l_{0}}-\rho_{\mathrm{lr}} \right)\left( \rho_{r_{0}}-\rho_{\mathrm{lr}} \right)-k_{\mathrm{off}}\left( f \right)\rho_{\mathrm{lr}},$$

where $\rho_{l_{0}}$ and $\rho_{r_{0}}$ are the total available ligand and receptor densities, respectively. Assuming that the bond turnover kinetics reach a quasi-steady state much faster than the cellular morphological changes, we can set $d\rho/dt\approx0$. This yields the equilibrium condition:

$$\begin{aligned} k_{\text{on}}\left( \rho_{l_{0}}-\rho_{\text{lr}} \right)\left( \rho_{r_{0}}-\rho_{\text{lr}} \right)=k_{\text{off}}\left( f \right)\rho_{\text{lr}}, \end{aligned}$$

where the dissociation rate constant, $k_{off}$, is force-dependent.

**Note B:** Geometric Model of Adherent Cells

The cell is modeled as a spherical cap. The effective surface area and volume read:

$$A_{\mathrm{eff}}=2\pi R\left( R+h \right),$$

$$V=\frac{1}{3}\pi\left( R+h \right)^{2}\left( 2R-h \right),$$

where $R$ represents the radius of the cell, while $h$ denotes the distance from the spherical center of the cellular dome to the adhesion plane. A positive $h$ value indicates the spherical center lies above the adhesion plane. Conversely, negative values indicate its position below this plane.

**Note C:** Ion Transport Mechanisms

In the process of osmotic shock, the ion transport serves as an essential part of the self-regulation of the cell. The ion transport consists of the passive ion transport and the active ion transport. The active transport could achieve ion flux against concentration gradients, whereas the passive transport only permits ion movements along concentration gradients. Under cellular steady-state conditions, the active and the passive transport achieve dynamic equilibrium. Mechanosensitive channel (MS channel) is a transmembrane protein, providing pathways for the passive ion transport. The activation of MS channels is related to the mechanical stress exerted on the membrane. This relationship can be approximately described as: When the mechanical stress on the membrane exceeds the critical stress threshold $\sigma_{c}$, MS channels start to open. Upon reaching the saturation stress $\sigma_{s}$, MS channels achieve maximal opening, with their gating extent becoming unresponsive to further stress increases. The ion flux through passive transport scales proportionally with both the opening degree of MS channels and their density on the membrane. Assuming the number of MS channels does not change during the process of osmotic shock, passive ion flux $J_{p}$ follows:

$$\begin{aligned} J_{passive}=\left\{ \begin{aligned} &0\text{ }\text{ }\text{if }\sigma\leq\sigma_{c} \\ &-\beta\left( \sigma-\sigma_{c} \right)\Delta\Pi\text{ }\text{if }\sigma_{c}<\sigma<\sigma_{s} \\ &-\beta\left( \sigma_{s}-\sigma_{c} \right)\Delta\Pi\text{ } \text{if }\sigma\geq\sigma_{s} \end{aligned} \right.,\#\left( 4 \right) \end{aligned}$$

in which $\beta$ is a rate constant.

The active transport relies on the active transmembrane protein which is directly related to ATP energy conversion. Cells exhibit an inherent upper limit in their capacity for active ion transport against concentration gradients. When osmotic pressure difference remains within critical osmotic pressure difference $\Delta\Pi_{c}$, cells can maintain active transport, with the ion transport rate exhibiting a negative correlation with the osmotic pressure difference. Contrarily, when $\Delta\Pi$ exceeds $\Delta\Pi_{c}$, active transporters fail to sustain against-gradient transport and instead facilitate passive ion movement along the concentration gradient. The active ion transport follows the relation:

$$\begin{aligned} J_{active}=\gamma\left( \Delta\Pi_{c}-\Delta\Pi\right),\#\left( 5 \right) \end{aligned}$$

$\gamma$ is a constant.

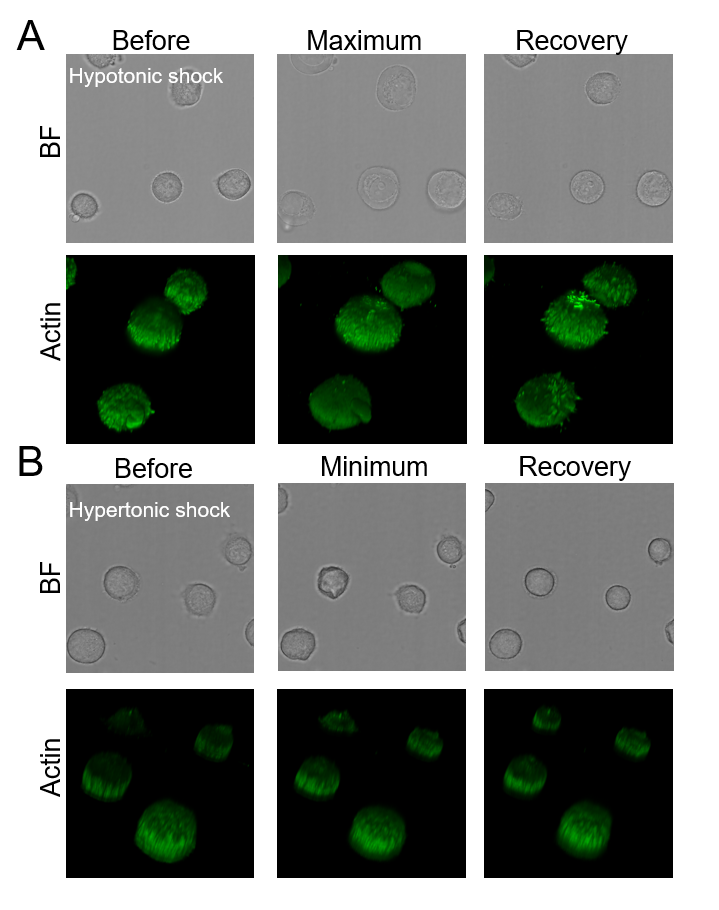

Fig. S1.

Validation of projected area as a proxy for cell volume. As shown in the figure, when the projected area increased, the volume revealed by three-dimensional reconstruction also increased; conversely, when the projected area decreased, the corresponding volume from 3D reconstruction decreased as well. (A) The expansion and recovery phases of the cell's bright-field projected area under hypotonic shock correspond well with the changes in three-dimensional volume. Fluorescence images are 3D reconstructions based on real-time imaging of actin-labeled cells, confirming that the volumetric changes align with variations in the projected area. (B) The correspondence between changes in cell volume and projected area under hypertonic stress.

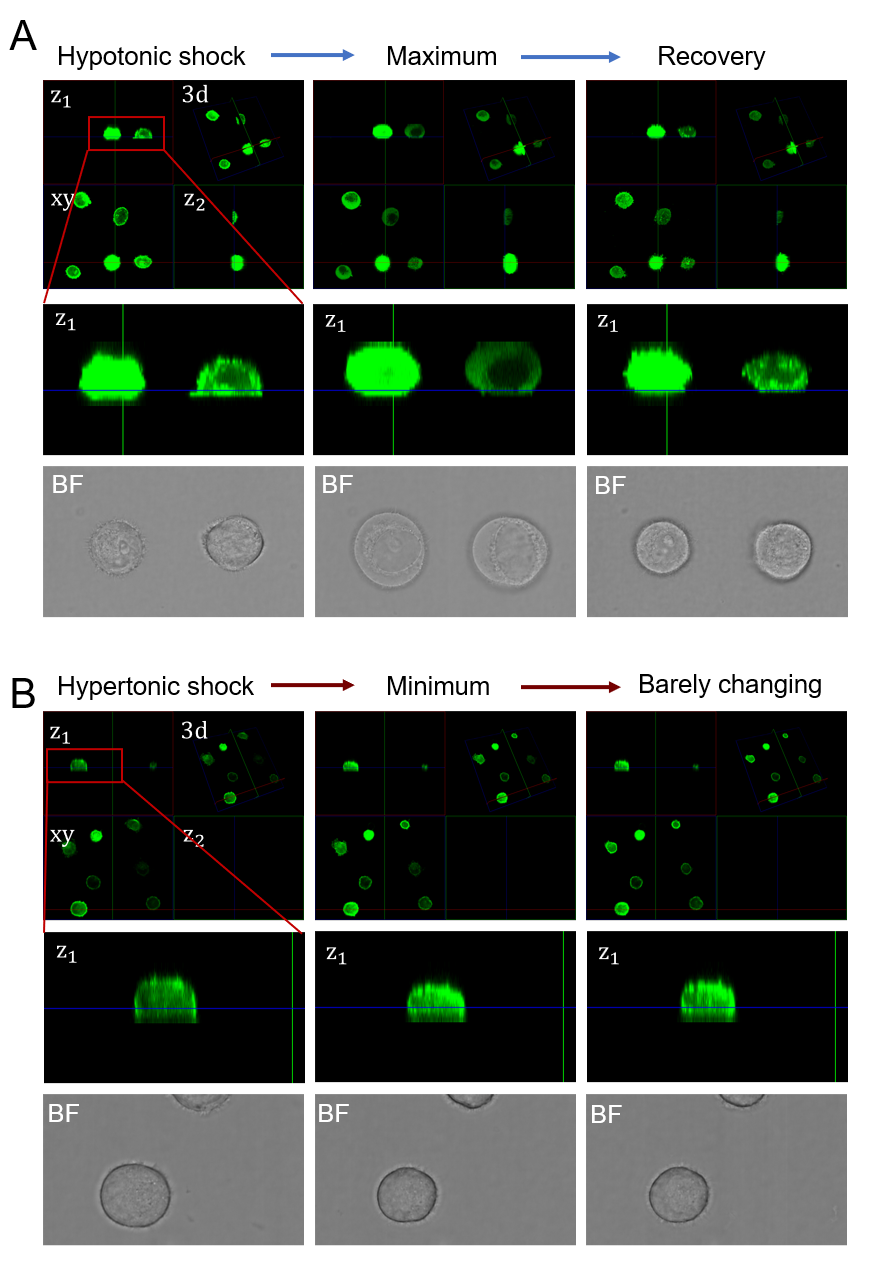

Fig. S2.

Z-axis height changes of cells under osmotic shock. As shown in the figure, an increase in projected area was accompanied by an increase in cell height along the z-axis, while a decrease in projected area corresponded to a reduction in z-axis height—further corroborating the accuracy of using projected area as a proxy for volume estimation. (A) Representative cell height changes along the z-axis under hypotonic shock. Panels z1 and z2 are longitudinal sections corresponding to the 3D rendering; xy denotes the top-down view. The fluorescence signal comes from labeled actin. (B) Representative cell height changes along the z-axis under hypertonic shock, with identical panel arrangement to (A).

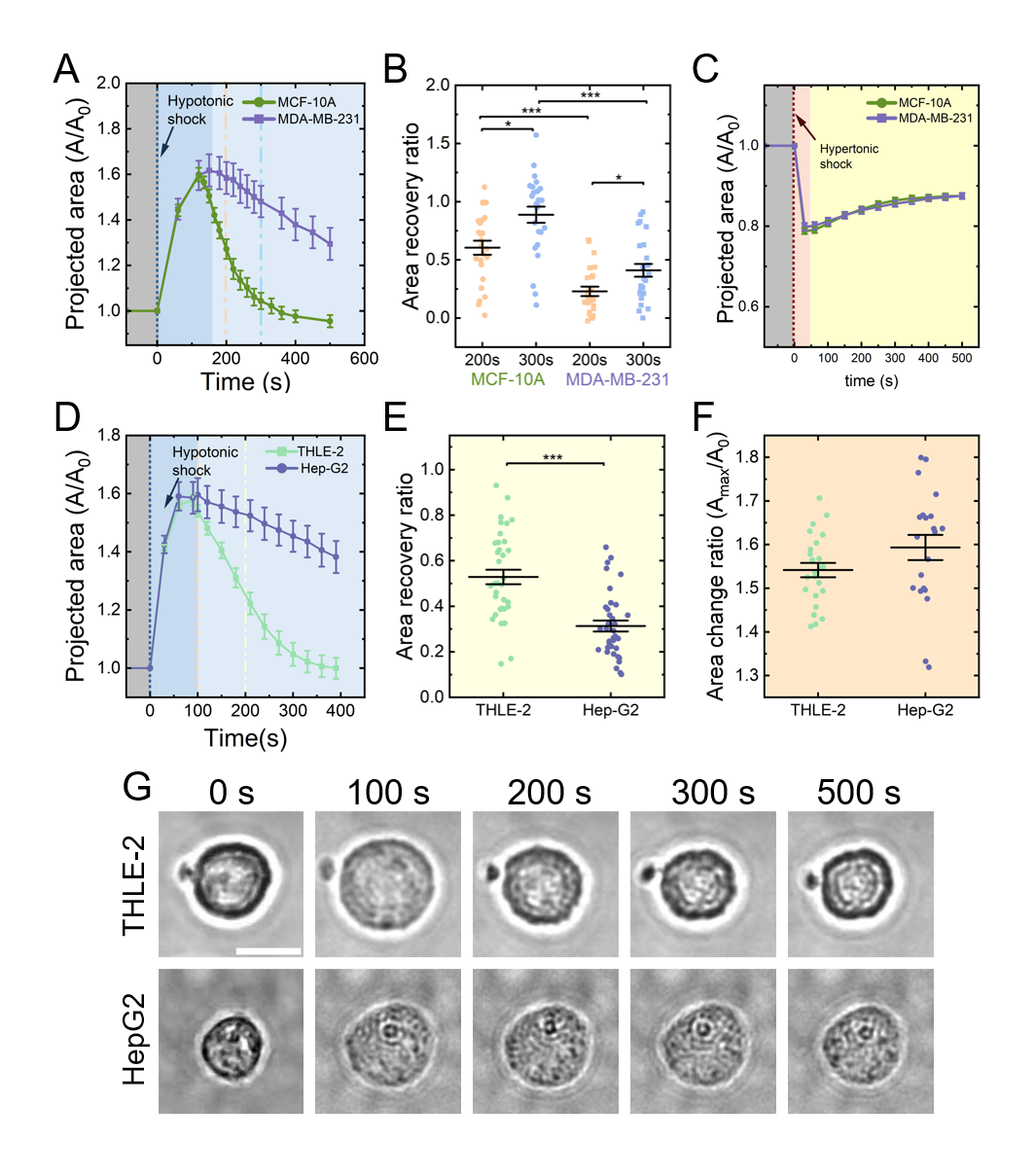

Fig. S3.

Volume responses to osmotic shocks across cell lines and conditions. (A) Normalized projected area dynamics of MCF-10A and MDA-MB-231 cells under hypotonic shock (200 mOsm). Data are mean ± SE (MCF-10A, n=14; MDA-MB-231, n=11). (B) Area recovery ratio at 200 s and 300 s after hypotonic shock (200 mOsm), calculated as (the maximum area - the recovery area) / (the maximum area – the initial area). Data are mean ± SE (MCF-10A, n = 25; MDA-MB-231, n = 25; mean ± SE). (C) Normalized projected area dynamics of MCF-10A and MDA-MB-231 cells under hypertonic shock (450 mOsm mannitol). Data are mean ± SE (MCF-10A, n=28; MDA-MB-231, n=29). (D) Normalized projected area dynamics of THLE-2 and HepG2 cells under hypotonic shock (200 mOsm). Data are mean ± SE (THLE-2, n=23; HepG2, n=25). (E) Area recovery ratio at 200 s after hypotonic shock (200 mOsm). Data are mean ± SE (THLE-2, n=24; HepG2, n=21). (F) Peak area change ratio (the maximum area / the initial area) at 100 s after hypotonic shock (200 mOsm). Data are mean ± SE (THLE-2, n=36; HepG2, n=36).

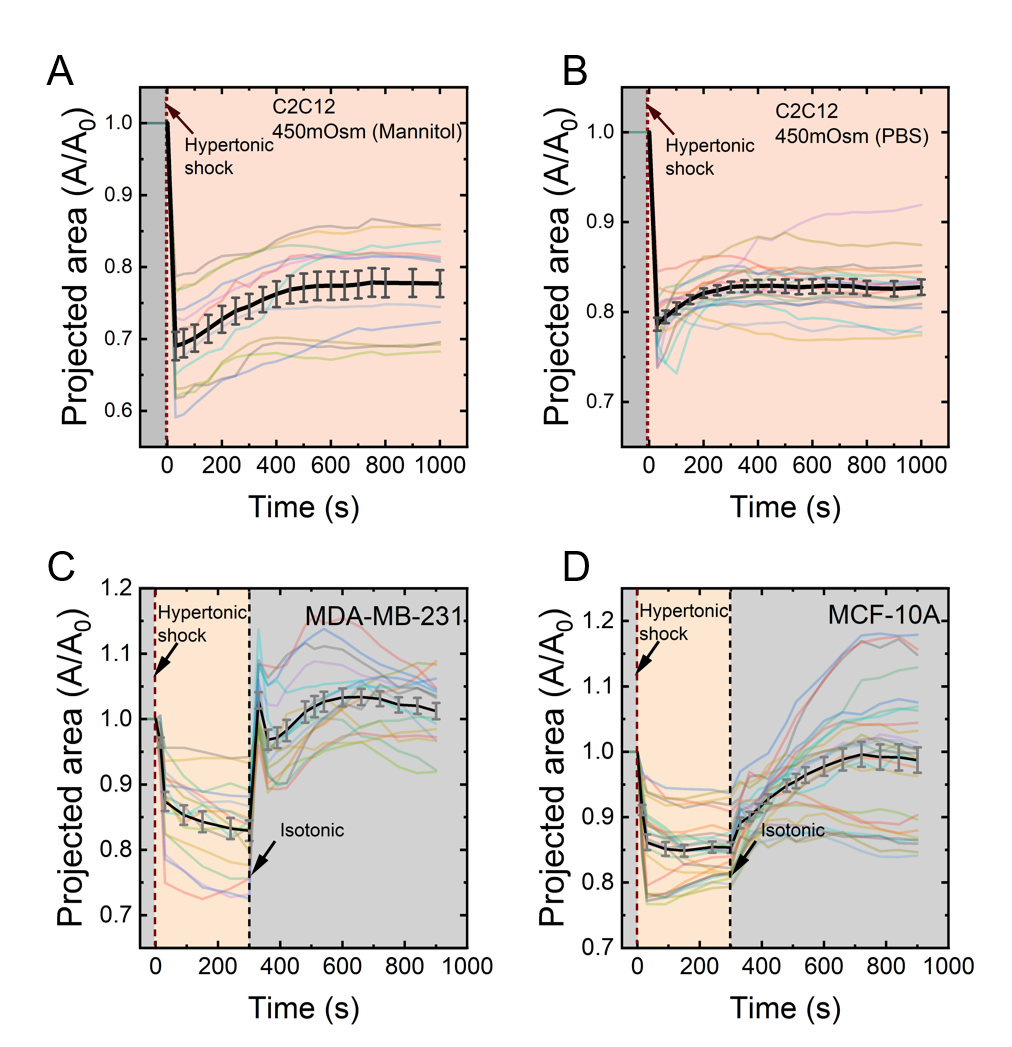

Fig. S4.

Volume responses of C2C12 and breast cells to hypertonic shock and recovery. (A) Normalized projected area dynamics of C2C12 cells under hypertonic shock (450 mOsm mannitol). Data are mean ± SE (n=12). (B) Normalized projected area dynamics of C2C12 cells under hypertonic shock (high-concentration PBS). Data are mean ± SE (n=17). (C) and (D) Normalized projected area dynamics of (C) MDA-MB-231 and (D) MCF-10A cells subjected to hypertonic shock (high-concentration PBS), with osmotic pressure returned to isosmotic levels at 300 s. Data are mean ± SE (MDA-MB-231, n=17; MCF-10A, n=27).

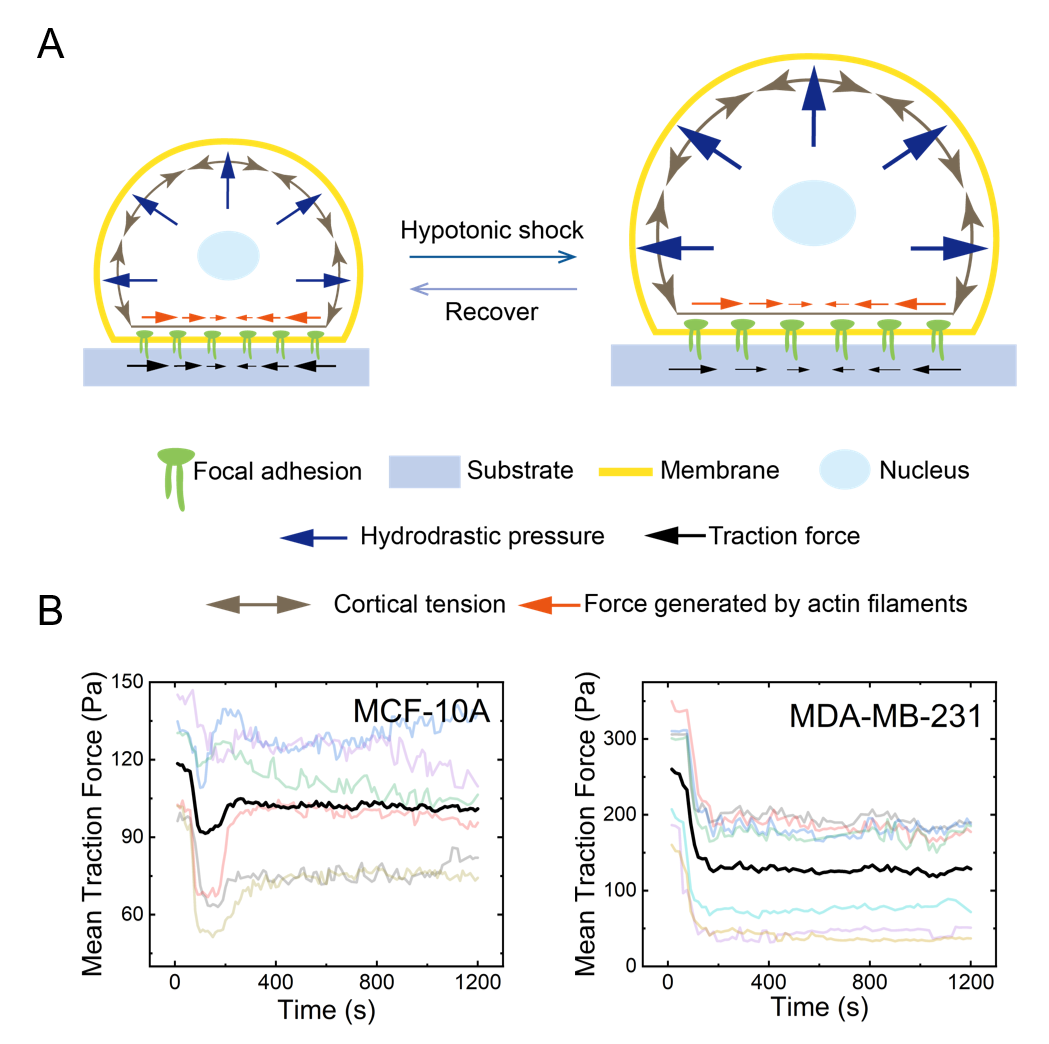

Fig. S5.

Traction force response to hypotonic shock. (A) Schematic of the mechanism underlying traction force changes induced by hypotonic shock. (B) Time course of the mean traction force for MCF-10A and MDA-MB-231 cells under hypotonic shock (100 mOsm). Data are mean (MCF-10A, n = 6; MDA-MB-231, n = 7).

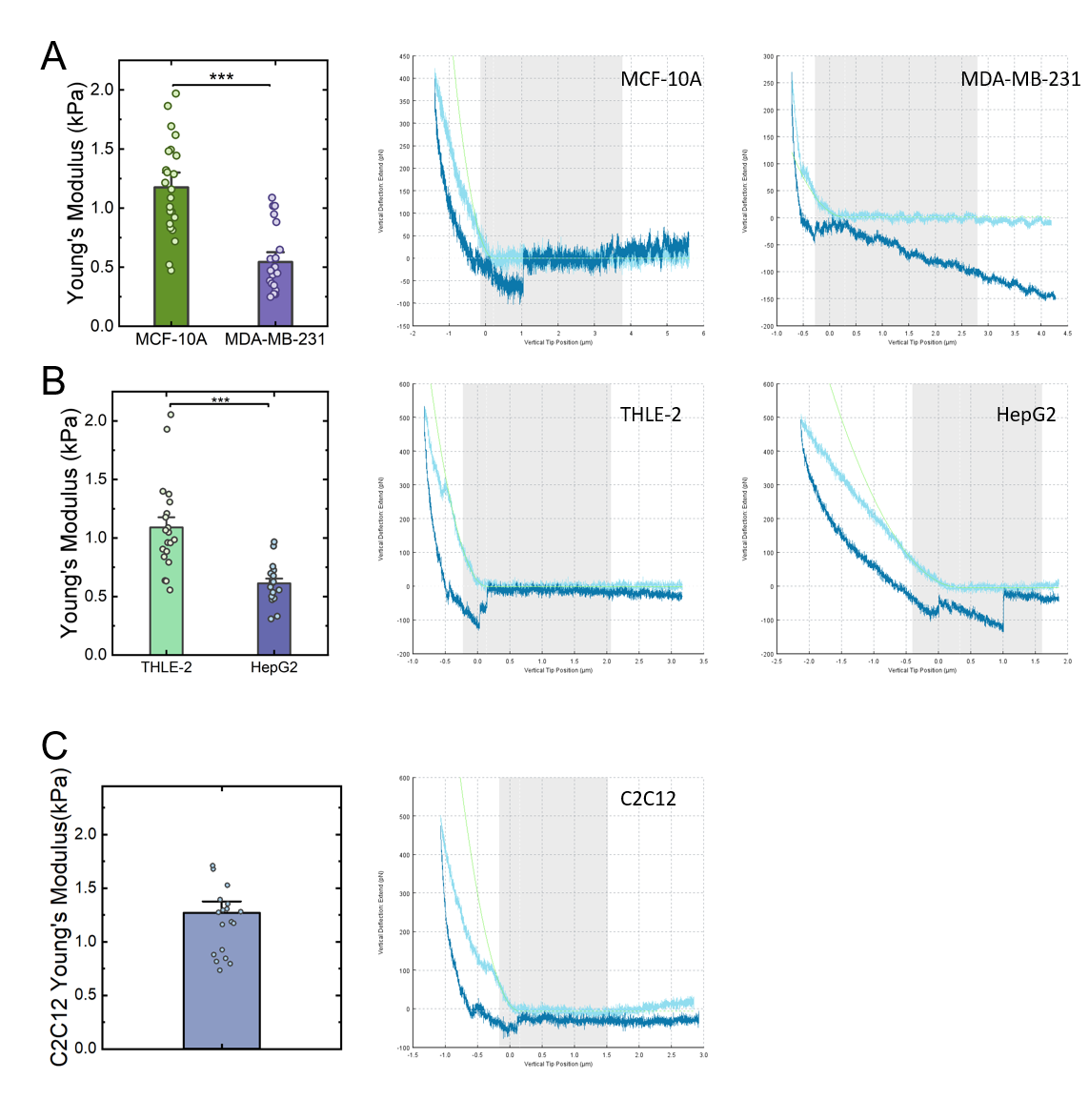

Fig. S6.

Cortical Young’s modulus measurements by atomic force microscopy (AFM). (A) Statistical analysis and representative AFM force-indentation curves for cortical Young's modulus of MDA-MB-231 and MCF-10A cells. Data are mean ± SE (MCF-10A, n=23; MDA-MB-231, n=23). (B) Statistical analysis and representative AFM data for cortical Young’s modulus of THLE-2 and HepG2 cells. Data are mean ± SE (THLE-2, n=20; Hep-G2, n=18). (C) Statistical analysis and representative AFM data for cortical Young's modulus of C2C12 cells. Data are mean ± SE (n=19).

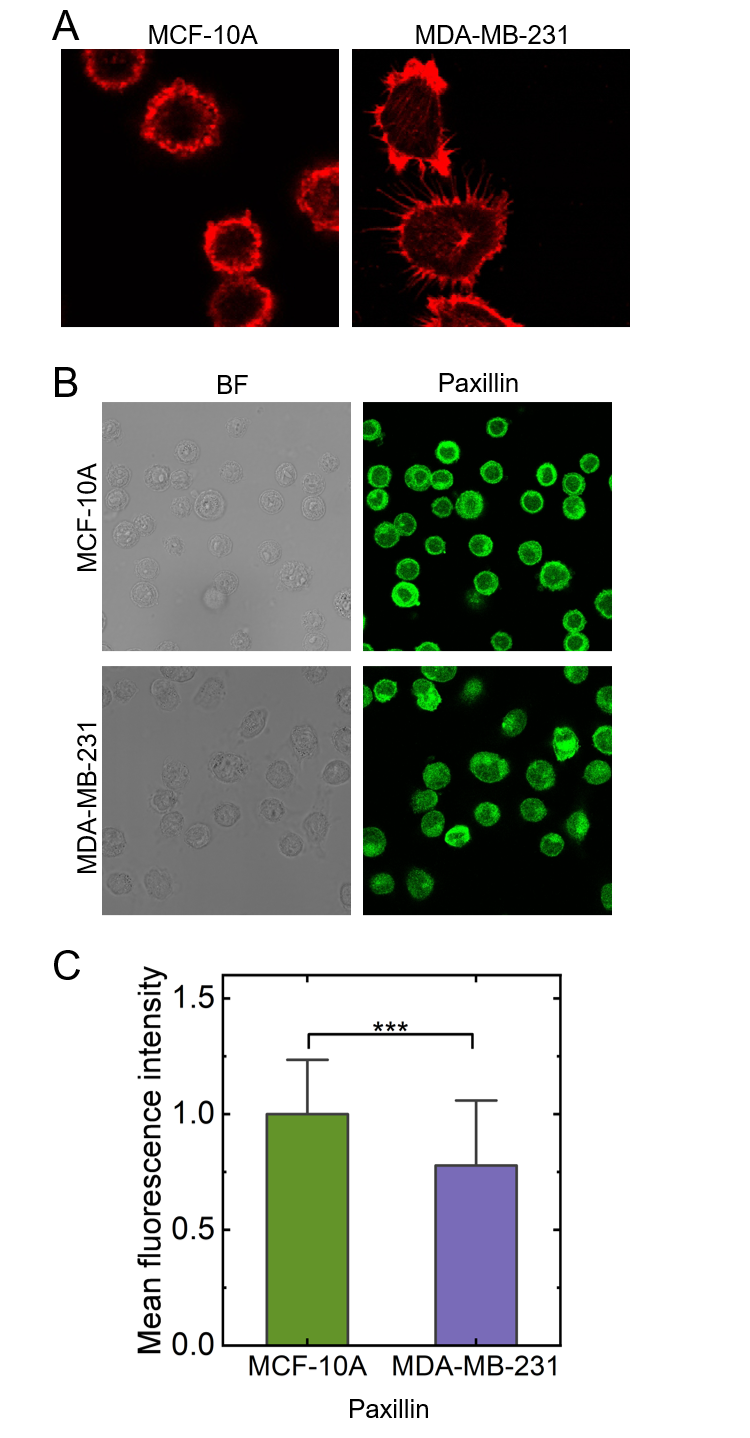

Fig. S7.

Characterization of basal actin structures and adhesion sites. (A) Representative fluorescence images of basal actin structures in slightly adherent MDA-MB-231 and MCF-10A cells, stained with phalloidin. (B) Representative bright-field and fluorescence images of paxillin in slightly adherent MDA-MB-231 and MCF-10A cells, visualized by immunofluorescence. (C) Quantification of the average fluorescence intensity from (B). Data are mean ± SE (MCF-10A, n=40; MDA-MB-231, n=39).

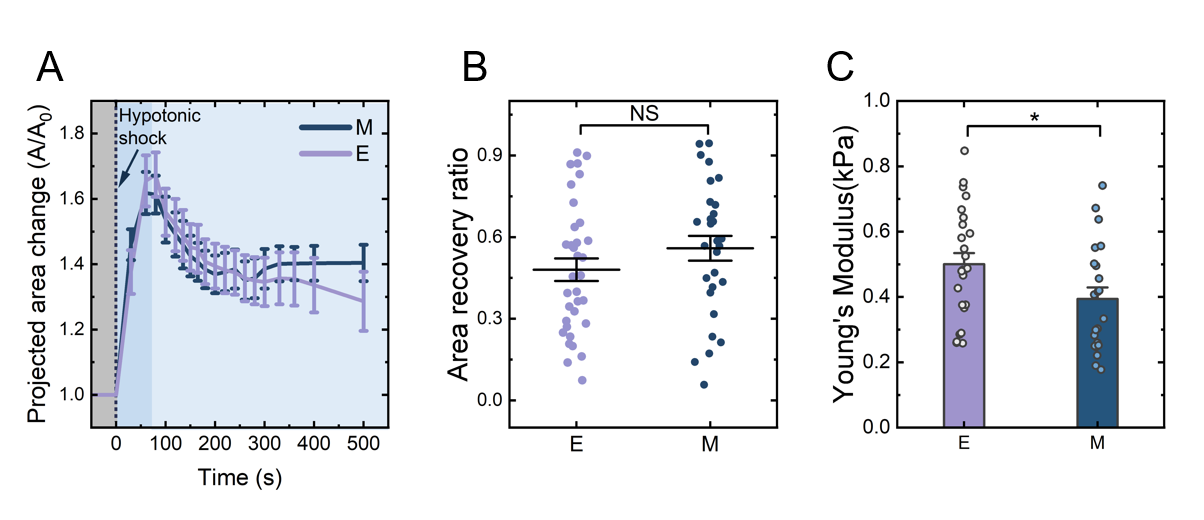

Fig. S8.

Phenotype-dependent volume response and cortical stiffness in HepG2 cells. (A) Normalized projected area dynamics of epithelial (E) and mesenchymal (M) phenotype HepG2 cells under hypotonic shock. Data are mean ± SE (E, n=10; M, n=9). (B) Area recovery ratio at 200 s after hypotonic shock (100 mOsm). Data are mean ± SE (E, n=34; M, n=29). (C) Cortical Young's modulus of E and M phenotype HepG2 cells. Data are mean ± SE (E, n=25; M, n=22).

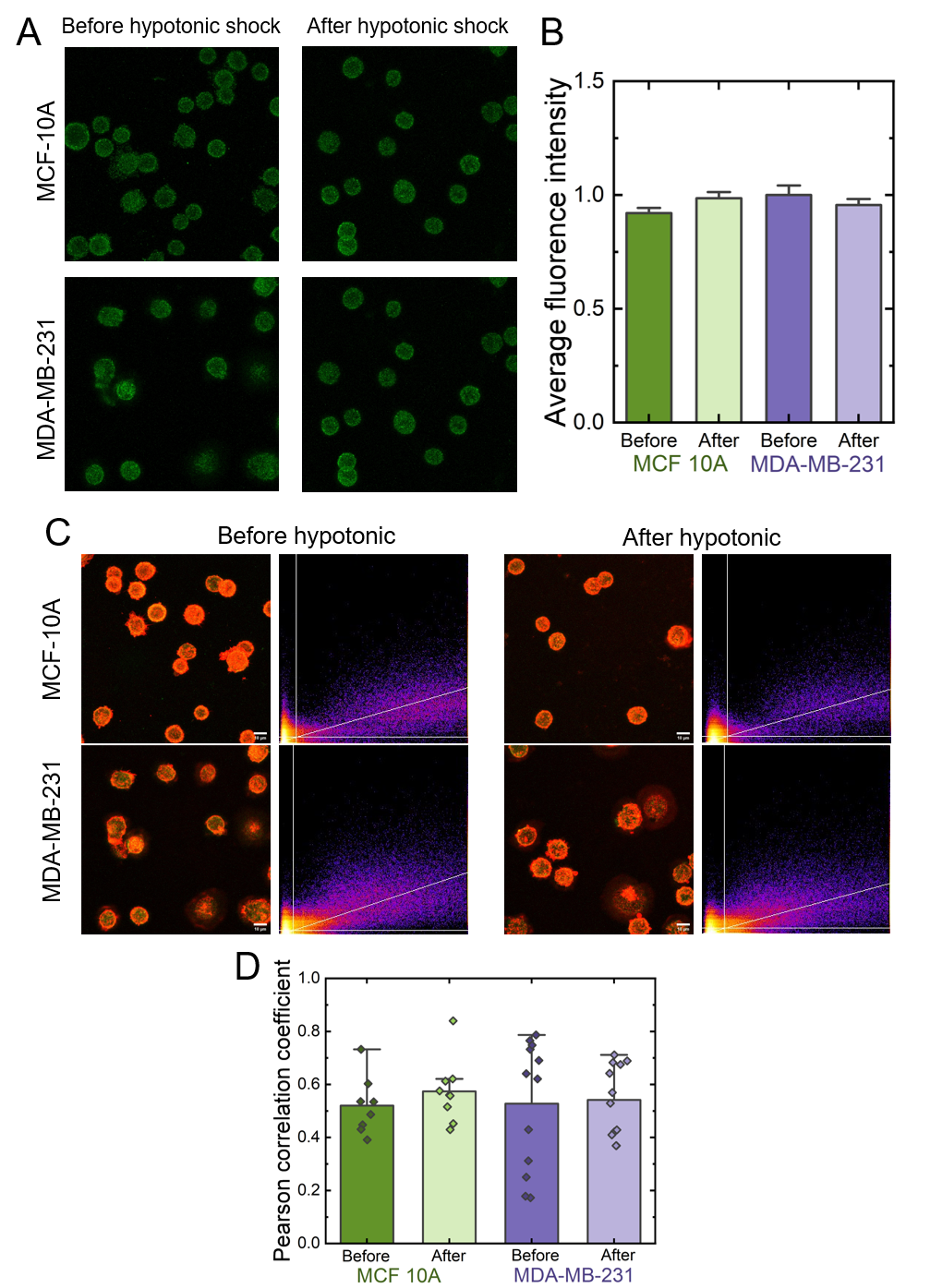

Fig. S9.

Myosin II distribution and co-localization with actin under osmotic shock. (A) Representative immunofluorescence images of myosin II before and after osmotic shock. (B) Quantification of the mean fluorescence intensity of myosin II. Data are mean ± SE (MCF-10A: before, n=23, after, n=21; MDA-MB-231: before, n=27, after, n=24). (C) Merged fluorescence images of actin (red) and myosin II (green) with co-localization analysis (yellow in merge). (D) Pearson's correlation coefficient for actin and myosin II co-localization before and after hypotonic shock. Data are mean ± SE (MCF-10A: before, n=8, after, n=8; MDA-MB-231: before, n=12, after, n=11).

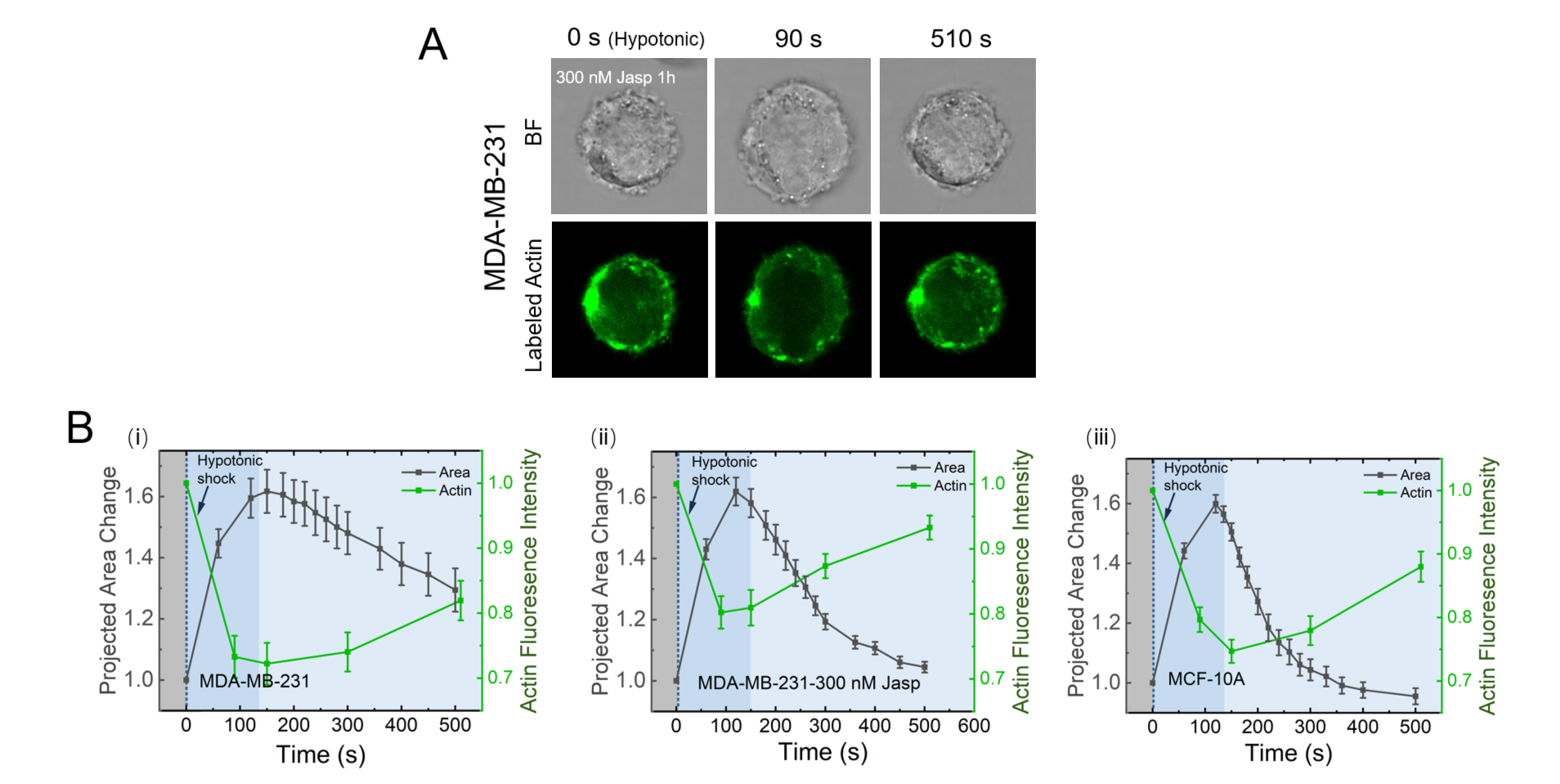

Fig. S10.

F-actin dynamics under hypotonic shock. (A) Representative bright-field and fluorescence images showing F-actin dynamics under hypotonic shock of MDA-MB-231 cells treated with 300 nM jasplakinolide for 1 h. (B) Time courses of the mean actin fluorescence intensity. (i) Untreated MDA-MB-231 cells. (ii) MDA-MB-231 cells treated with 300 nM jasplakinolide for 1 h. (iii) MCF-10A cells. Data are mean ± SE (Untreated MDA-MB-231, n=12; Jasplakinolide-treated MDA-MB-231, n=15; MCF-10A, n=18).

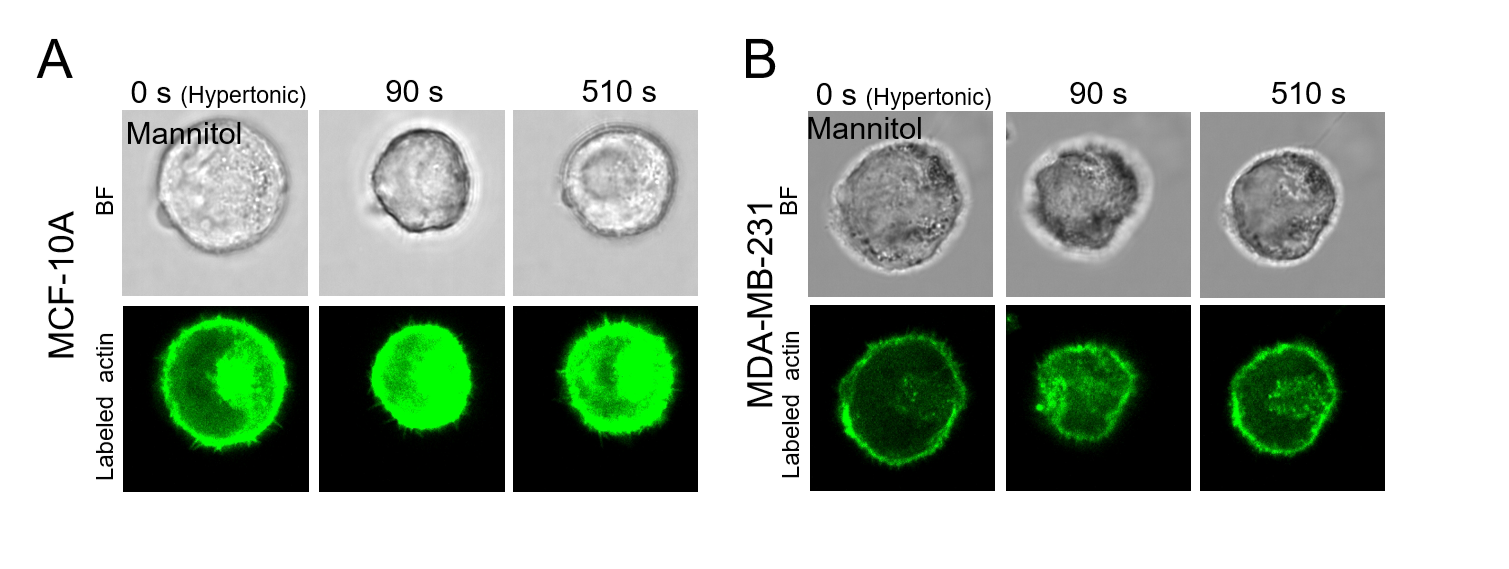

Fig. S11.

F-actin dynamics under hypertonic shock. (A) Representative images of F-actin dynamics in MCF-10A cells under hypertonic shock induced by 450 mOsm mannitol. (B) Representative images of F-actin dynamics in MDA-MB-231 cells under hypertonic shock induced by mannitol.

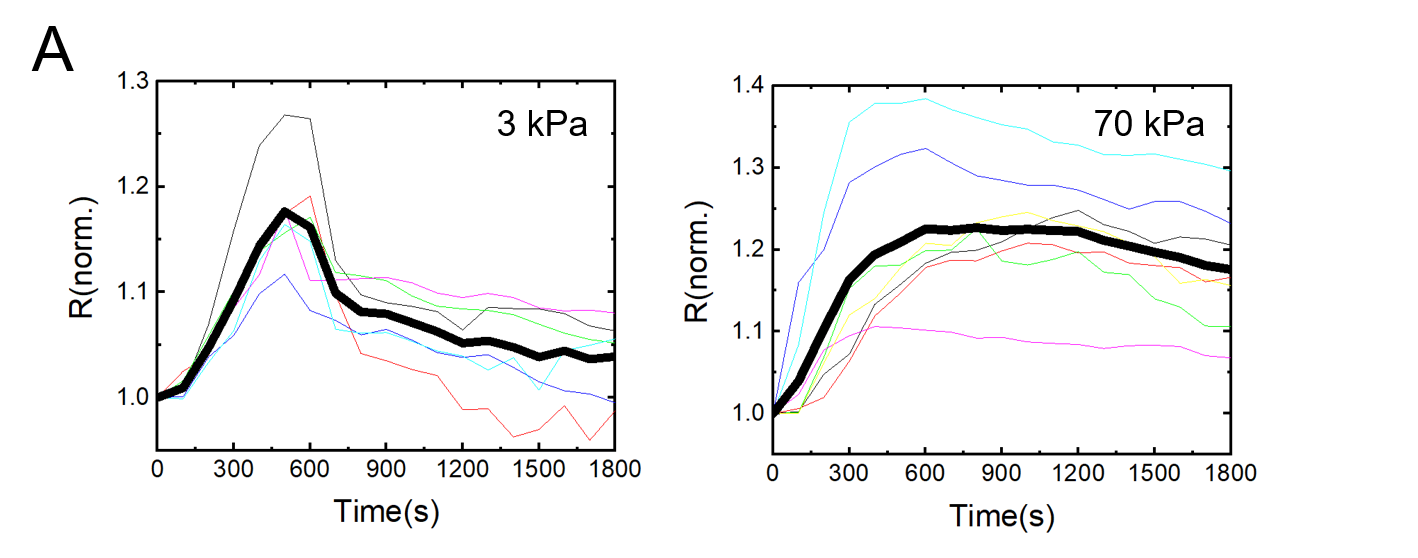

Fig. S12.

Effect of substrate stiffness on volume recovery in cancer cells. Normalized cell radius dynamics of MDA-MB-231 cells under hypotonic shock on substrates with stiffnesses of 3 kPa and 70 kPa.

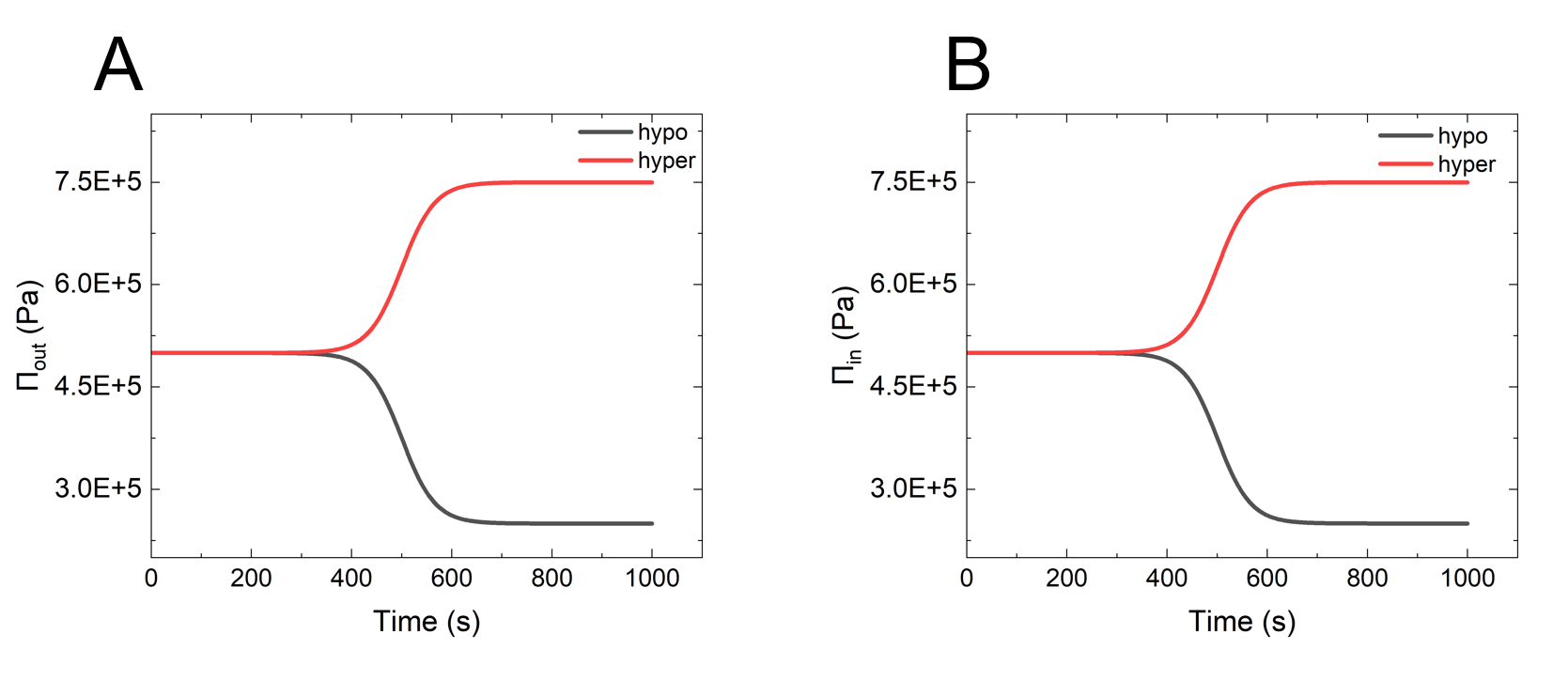

Fig. S13.

Simulated osmotic pressure changes: (A) extracellular osmotic pressure; (B) intracellular osmotic pressure.
